# *Ex vivo* and *in vivo* evidence that cigarette smoke-exposed T regulatory cells impair host immunity against *Mycobacterium tuberculosis*

**DOI:** 10.1101/2022.01.20.477097

**Authors:** Xiyuan Bai, Deepshikha Verma, Cindy Garcia, Ariel Musheyev, Kevin Kim, Lorelenn Fornis, David E. Griffith, Li Li, Nicholas Whittel, Jacob Gadwa, Tamara Ohanjanyan, Diane Ordway, Edward D. Chan

## Abstract

A strong epidemiologic link exists between exposure to cigarette smoke (CS) and increased susceptibility to tuberculosis (TB). *In vitro* macrophage and *in vivo* murine studies showed that CS and nicotine impair host-protective immune cells against *Mycobacterium tuberculosis* (*MTB*) infection. However, little is known about how CS may affect immunosuppressive cells in the context of *MTB* infection. Thus, we investigated whether CS-exposed T regulatory cells (Tregs) could exacerbate *MTB* infection in co-culture with human macrophages and in the adoptive transfer of Tregs from air- and CS-exposed mice. We found that exposure of primary human Tregs to CS extract impaired the ability of human monocyte-derived macrophages to control an *MTB* infection by inhibiting phagosome-lysosome fusion and autophagosome formation. Neutralization of CTLA-4 on the CS extract-exposed Tregs abrogated the impaired control of *MTB* infection in macrophage and Treg co-cultures. In Foxp3^+^GFP^+^DTR^+^ (Thy1.2) mice depleted of endogenous Tregs, adoptive transfer of Tregs from donor CS-exposed B6.PL(Thy1.1) mice with subsequent *MTB* infection of the recipient Thy1.2 mice resulted in a greater burden of *MTB* in the lungs and spleens than those that received Tregs from air-exposed mice. Mice that received Tregs from CS-exposed mice and then infected with *MTB* had modest but significantly reduced numbers of interleukin-12-positive dendritic cells and interferon-gamma-positive CD4^+^ T cells in the lungs and increased number of programmed cell death protein-1 positive CD4^+^ T cells in both the lungs and spleens.

## INTRODUCTION

Worldwide, there are marked geographic overlaps of cigarette smoking and tuberculosis (TB) cases (1). Epidemiological studies show that individuals with cigarette smoke (CS) exposure have increased link to increased: infection rate, progression of primary infection, reactivation TB, severe disease, delay in sputum conversion, recurrence (relapse), worse treatment outcome, and TB mortality (2–4). While CS has deleterious effects on anti-TB immunity (5–10), the impact of CS on immunoregulating cells, particularly T regulatory cells (Tregs) has been less well studied (11). O’Leary *et al* (7) noted that both smokers’ and nonsmokers’ alveolar macrophages could induce phenotypic expression of Tregs.

Murine studies have shown that knockdown of Tregs in the *early* phase of *Mycobacterium tuberculosis* (*MTB*) infection results in reduced *MTB* burden, indicating that excessive number of Tregs soon after infection impairs host immunity against *MTB* (12). In contrast, Tregs play a less important role in later stages of infection (12). Moreover, active TB patients have higher Treg numbers than those with latent TB infection (13). Thus, a vigorous early influx of activated Tregs to the lungs, particularly before *MTB* infection is under control, worsens TB in experimental animals and as supported by congruent findings in primary human cells (14–19).

Previously, our group and others showed that CS and nicotine could impair the ability of macrophages to control an *MTB* infection (5–11). However, since CS and nicotine may (*i*) increase the influx and function of Tregs in the lungs (20–23), (*ii*) significantly increase the number of alveolar macrophages (24), and (*iii*) alveolar macrophages of smokers can drive naïve T cells toward the Treg phenotype (7), it is plausible that CS could also negatively impact host immunity against *MTB* indirectly by enhancing Treg activity. To further elucidate the effects of CS on anti-TB host immunity, we undertook an *ex vivo* study of co-culture of human macrophages and air- or CS extract-exposed Tregs as well as an *in vivo* murine study using adoptive transfer of Tregs from air- or CS-exposed donor mice to determine whether CS-exposed Tregs could exacerbate *MTB* infection.

## EXPERIMENTAL METHODS

### *Mycobacterium tuberculosis* strains, mice, and reagents

H37Rv-*MTB* was obtained from American Type Culture Collection (ATCC, Manassas, VA). HN878 W-Beijing strain of *MTB* (HN878-W-Beijing *MTB*) was originally a kind gift from Dr. B. Kreiswirth (Public Health Research Institute Center, Newark, NJ). The B6.PL(Thy1.1) and Foxp3^+^GFP^+^DTR^+^ (Thy1.2) mice were purchased from Jackson Laboratories, Bar Harbor, Maine. Cell Preparation Tubes (CPT) used to isolate peripheral blood mononuclear cells (PBMC) were purchased from BD Company (San Jose, CA). The Miltenyi Biotech Regulatory T Cell Isolation Kit II (human) (Auburn, CA) was used to isolate human Tregs from peripheral blood. 4’,6-diamidino-2-phenylindole (DAPI) and LysoTracker Red DND-99 were purchased from Invitrogen, Carlsbad, CA. Macrophage colony-stimulating factor (M-CSF, human) was purchased from Millipore Sigma, St. Louis, MO. Fetal bovine serum was acquired from Atlanta Biologicals (Norceross, GA) and inactivated at 56°C for one hour. The LC3-IIB and p62 primary rabbit antibodies were purchased from Cell Signaling Technology Inc. (Danvers, MA). Roswell Park Memorial Institute (RPMI) medium, CY3 goat anti-rabbit IgG (H+L) Cross-Adsorbed Secondary Antibody, and the four and eight chambered Nunc^™^ Lab-Tek^™^ II Chamber Slide^™^ System were purchased from ThermoFisher Scientific (Waltham, MA). PE-conjugated anti-CTLA-4 and APC-conjugated anti-CD279 (PD-1) were purchased from R&D System (Minneapolis, MN). Human cytokines from cell culture supernatants were analyzed using ELISA kits for TNF, IL-10, and TGFβ (R&D Systems, Minneapolis, MN). Anti-CTLA-4 neutralizing antibody and non-immune human IgG antibody were purchased from BPS Bioscience Inc (San Diego, CA). Diphtheria toxin was purchased from List Biological Laboratories, Campbell, CA. Information on other reagents are discussed in the relevant methods sections below.

### Cigarette smoke (CS) extract preparation

CS extract was prepared by mechanically “smoking” a single unfiltered 3R4F cigarette into 10 mL of pre-warmed RPMI medium via a vacuum pump, with the flow adjusted at 3 L/min using an Accucal flowmeter (Gilmont Instruments). The extract was then filtered through a 0.22 μm syringe filter and adjusted to a pH of 7.4. This solution is arbitrarily designated as 100% CS extract (25).

### Blood processing and Treg isolation

After informed consent (Colorado Multiple Institutional Board Review, protocol 16-1413), blood was drawn into six Cell Preparation Tubes^®^ from each healthy donor. Peripheral blood mononuclear cells (PBMC) were isolated by centrifugation. Following the determination of the total cell count, half of the PBMC aliquot was cryopreserved in freeze medium (85% FBS, 15% DMSO) for future Treg isolation. The other half was incubated with 20 ng/mL of monocyte-colony stimulating factor for one week to prepare monocyte-derived macrophages (MDM).

One day before completion of MDM differentiation, an autologous cryopreserved PBMC aliquot was thawed and then centrifuged to remove the DMSO. Tregs were isolated from PBMC using the Regulatory T Cell Isolation Kit II according to manufacturer’s instructions. In brief, non-CD4^+^ and CD127^high^ cells were magnetically labeled and then depleted. From the remaining cells, CD25^+^ cells were magnetically labeled and then isolated, yielding only cells with biomarkers CD4^+^CD25^+^CD127^dim/-^, the cell surface markers of Tregs.

### MDM infection and co-culture to quantify CFU

Differentiated MDM were infected with *MTB* H37Rv at a multiplicity-of-infection of 10 *MTB:1* macrophage for one hour, and then washed gently to remove any free *MTB*. MDM were then incubated with: (*i*) RPMI medium + 10% FBS alone or with (*ii*) non-conditioned Tregs or (*iii*) Tregs previously conditioned in 5% CS extract for 18 to 24 hours. After an additional hour of incubation, Day 0 cells were washed, lysed, serially diluted, and plated on 7H10 agar to quantify *MTB*. The remaining cells were incubated for 2 or 4 days before quantifying *MTB*.

### Phagosome-lysosome fusion

PBMC were differentiated into MDM in four-chamber glass slides. After one week, the MDM were co-cultured with autologous Tregs (100 MDM:1 Treg ratio) that had been previously cultured in medium alone or with 5% CS extract for 18 to 24 hours. MDM alone and MDM co-cultured with air- or CSE-exposed Tregs were infected with GFP-*MTB* H37Rv at a MOI of 10 *MTB:1* macrophage for six hours. At the 4-hour mark, 25 μL of LysoTracker Red (50 nM) was added to each chamber well containing 500 μL of medium to label the lysosomes. At the 6-hour time point, the medium was removed, the cells fixed with 4% paraformaldehyde, washed thrice with 1X PBS, stained with DAPI and stored overnight to dry in the dark at room temperature. The following day, the cells were viewed under 40X oil immersion lens with an inverted Zeiss 200M microscope and a 175-watt xenon lamp in a DG4 Sutter instrument lamp housing. The Cy3 filter was used to detect the LysoTracker-stained lysosomes while FITC was used to detect GFP-*MTB*. Co-localization of the GFP-*MTB* with lysosomes would appear yellow. Ten random pictures were taken per well with Live-Cell Marianis. Images were further analyzed using FIJI. P-L fusion was calculated by dividing the number of cells with evidence of GFP-*MTB* and lysosome fusion by the total number of cells containing GFP-*MTB* with or without GFP-*MTB*-lysosome fusion.

### Autophagosome formation and maturation

MDM were co-cultured in four-chamber glass slides with Tregs (100 MDM:1 Treg) that had previously been exposed to medium alone or 5% CS extract for 18 to 24 hours. After one hour of incubation with Tregs, the cells were infected with GFP-*MTB* (MOI of 10 *MTB:1* macrophage) and incubated for 18 hours at 37°C and 5% ambient CO_2_. The cells were then washed with PBS, fixed with 4% paraformaldehyde for 30 minutes, washed, incubated with permeabilization buffer (0.5% Triton X in PBS) for 10 minutes, rinsed with PBS, and incubated with blocking buffer (5% BSA, 0.5% Tween 20 in PBS) for one hour. Anti-LC3 antibody at a dilution of 1.5:200 was incubated with the cells and left in a dark container at 4°C overnight. An anti-rabbit CY3 tagged antibody (1:1000) was added to the wells, left in the dark at room temperature for 45 minutes, and the cells fixed with DAPI. The slides were dried overnight and viewed with the Live-Cell Marianis microscope to analyze and quantify autophagosome formation. Multiple images were taken from all conditions and five were picked at random. The number of LC3-II puncta per cell was determined and averaged for 15 randomly chosen cells. The same process was replicated for all three conditions.

To determine autophagosome maturation, we quantified p62-positive puncta in a temporal fashion of MDM, MDM + unexposed Tregs, and MDM + CS extract-exposed Tregs that were infected with GFP-*MTB* for 3, 6, and 18 hours. At the end of each time point, the cells were washed with 1X PBS and fixed with 4% paraformaldehyde for 30 minutes. The cells were washed, permeabilized for 10 minutes, rinsed with 1X PBS, incubated with blocking buffer for one hour. Lastly, 25 μL of anti-p62-rabbit antibody [1:200] was added to each chamber well and incubated overnight at 4°C. Then CY3 anti-rabbit [1:1000] secondary antibody was added and incubated for 45 minutes in the dark. After incubation, slides were rinsed, DAPI added, and once dried, the cells were viewed under a 40X oil emersion lens. CY3 filter was used for detecting p62 and FITC for GFP-*MTB*. Further analyses of p62 in the presence of 200 nM rapamycin and 100 nM bafilomycin were conducted similarly.

### CS exposure of mice

The B6.PL(Thy1.1) mice were exposed to CS in a whole animal exposure chamber using smoke generated by the TE-10c cigarette smoking machine (Teague Enterprises) as previously described (9). The relative proportion of smoke particulate, vapors, and gases generated approximates environmental CS exposure (89% side-stream smoke exposure and 11% mainstream CS exposure), maintaining Total Suspended Particulate concentration of 85-120 mg/m^3^ for 5 hours a day and 5 days a week for four weeks.

### Depleting, harvesting, and adoptively transferring Tregs

Ten days before the adoptive transfer of Tregs, endogenous Tregs of the recipient Foxp3^+^GFP^+^DTR^+^ (Thy1.2) mice were depleted *via* administration of diphtheria toxin at 50 μg/kg x four doses. Tregs of the air- and CS-exposed B6.PL(Thy1.1) mice were harvested by cell sorting. To sort CD4^+^CD25^hi^CD127^-^ and CD4^+^CD25^-^ cells, single cell suspension was prepared from the lungs of donor B6.PL(Thy 1.1) mice. After incubating with 5 μg/mL Fc block (ebioscience) for 20 min at 4°C, the cells were attained with APC labeled anti-CD4 (clone gk1.1; ebioscience), Pe Cy7-labeled anti-CD25 (clone PC61.5; ebioscience), Alexa ef450 labeled CD127 (eBioRDR5; ebioscience) and FITC-labeled anti-CD3 (clone 17A2; ebioscience). Thereafter, the cells were stained with 0.5 μg/ml 7-AAD (eBioscience) for 5 min, sorted on a FACS Aria III (BD Biosciences) with a 70-μm nozzle, doublets gated out by FSC-A vs. FSC-H, and 7-AAD^+^ dead cells excluded from the singlet population using FL3. The live CD4^+^CD25^hi^CD127^-^ or CD4^+^CD25^-^ were individually sorted. One million donor Tregs were transferred to each Treg-depleted Foxp3^+^GFP^+^DTR^+^ (Thy1.2) mouse *via* intravenous tail vein injection.

### Infection of Foxp3^+^GFP^+^DTR^+^ (Thy1.2) mice with *MTB*

Once the Foxp3^+^GFP^+^DTR^+^ (Thy1.2) mice received the Tregs from the donor B6.PL(Thy1.1) mice, the recipient mice were infected with HN878 W-Beijing strain of *MTB* using the Glas-Col Aerosol Generator at an inoculum dose of 2×10^6^ CFU/mL. The mice were exposed to an aerosol infection in which approximately 100 bacteria were deposited in the lungs of each mouse.

### *MTB* enumeration in mice

One, 30 and 60 days after infection, the mice were sacrificed by CO_2_ inhalation, and their lungs and spleen were prepared to quantify *MTB* (CFU) and immune cell phenotypes by flow cytometry and analyze the lungs by histopathology. The lungs and spleens were homogenized in saline and serial dilutions were plated on 7H11 agar plates supplemented with OADC (BD Biosciences, San Jose, CA). After 3-4 weeks of incubation at 37°C, CFUs were counted and the data expressed as the log_10_ numbers per target organ.

### Flow cytometry for surface markers and intracellular cytokines

For flow cytometry analysis, single-cell suspensions of lungs and spleen from each mouse were resuspended in PBS containing 0.1% sodium azide. Fc receptors were blocked with purified anti-mouse CD16/32. The cells were incubated in the dark for 25 minutes at 4°C with predetermined optimal titrations of specific antibodies. Cell surface expression was analyzed for CD279 (PD-1) (clone J43), CD4 (clone GK1.5), CD11c (clone HL3), and CD11b (clone M1/70). All antibodies and reagents were purchased from BD Pharmingen (San Diego, CA). The samples were probed using a Becton Dickinson FACS canto instrument, and the data were analyzed using FlowJo software. Individual cell populations were identified according to the presence of specific fluorescence-labeled antibodies. All the analyses were performed with the acquisition of a minimum of 300,000 events.

### Intracellular cytokine staining

The cells were initially stimulated for 4 hours at 37°C with a 1X cell stimulation cocktail (eBioscience) diluted in complete DMEM. Thereafter, the cells were stained for cell surface markers, as indicated, fixed, and permeabilized according to the manufacturer’s instructions for the Fix/Perm and Perm wash kit (eBioscience). The cells were then incubated for 30 minutes at 4°C with FcBlock plus anti–IL-10 (clone JES5-16E3; eBioscience), anti–IFNγ (clone XMG1.2; eBioscience), anti-IL-17, anti-Foxp3 (clones FJK-16s; eBioscience), anti-IL-12 (clone C15.6; BD), anti-TNF (clone MP6-XT22; eBioscience), anti-CTLA-4 (clone 63828; R&D Systems) or with the respective isotype control. Data acquisition was performed on FACS canto cytometer (BD) and analysis was performed as described (26).

## RESULTS

### Confirmation of Treg isolation

Tregs were isolated from PBMC using the Miltenyi Biotech Regulatory T Cell Isolation Kit II. After staining with CD127-BB515, CD25-PE, and CD4-APC-CY7, the cells were analyzed with LSR Fortessa equipped with FACSDiva (BD Biosciences) and then with FlowJo software. As shown in **Supp Figure 1**, essentially all the isolated cells had markers entirely consistent with Tregs.

### CS extract-exposed Tregs impaired macrophage control of *MTB* infection

To determine whether CS extract-exposed human Tregs impair macrophage control of *MTB*, MDM were first infected with *MTB* for 1 hour, washed to removed free *MTB*, and then cocultured with autologous primary human Tregs (at a ratio of 100 monocytes: 1 Treg) that were previously incubated in medium alone or with 5% CS extract for 18-24 hours. It is important to note that the Tregs were washed to remove any medium containing CS extract before coculture with the MDM to ensure that any effect of CS extract was caused by the CS extract-exposed Tregs and not due to any direct effects of CS extract on MDM. After the *MTB*-infected cell co-cultures were incubated for 1 hour, 2 and 4 days, cell-associated *MTB* were quantified as previously reported (11). As an additional control, MDM without Tregs were also infected with *MTB*. Compared to MDM alone, co-culture of MDM with control Tregs increased the bacterial burden, further increasing in co-culture with CS extract-exposed Tregs (**Figure 1A**).

**Figure 1.**
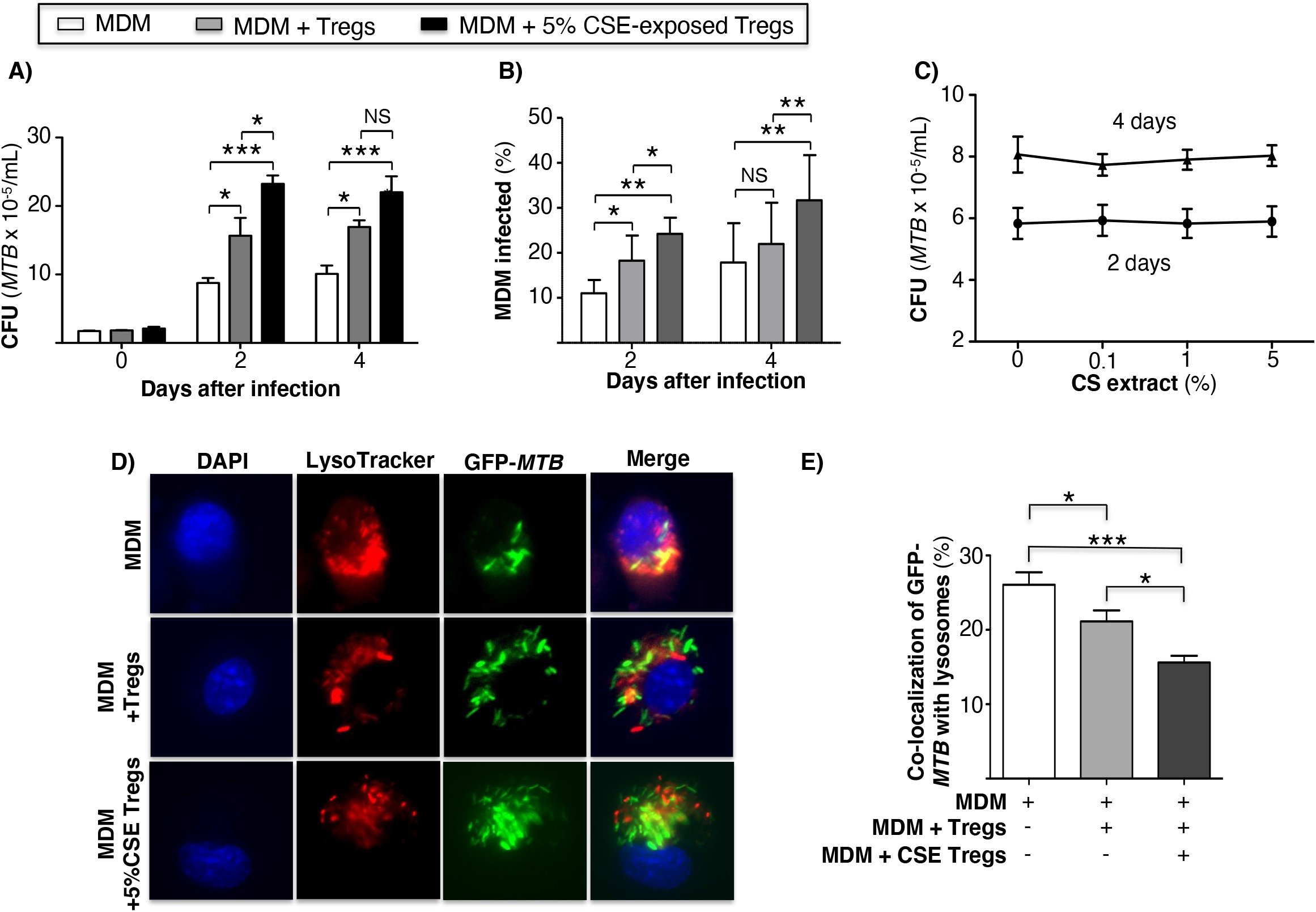
Cigarette smoke extract-exposed T regulatory cells impairs macrophage control of *Mycobacterium tuberculosis* infection by reducing phagosome-lysosome fusion. **(A)** *MTB*-infected human monocyte-derived macrophages (MDM) alone or co-incubated with unexposed or 5% CS extract-exposed Tregs for the indicated times and intracellular *MTB* (CFU) quantified. **(B)** MDM were infected with GFP-*MTB* for 1 hour, washed, incubated with medium alone or co-incubated with unexposed or 5% CS extract-exposed Tregs, and at the indicated times, the cells were prepared for fluorescent microscopy. The percent of cells infected was calculated by dividing the number of GFP-*MTB* infected MDM by the total number of at least 500 MDM cells counted per condition. Data in (A) and (B) represent the mean ± SEM of experiments of cells performed in duplicates from three subjects. **(C)** *MTB* were cultured with 7H9 medium alone or with the indicated concentrations of CS extract for 2 and 4 days and *MTB* was quantified. **(D)** MDM were incubated in medium alone or co-cultured with unexposed or 5% CS extract-exposed Tregs, followed by infection with *GFP-MTB* and assayed for P-L fusion. Shown are representative images of the identified conditions. **(E)** P-L fusion percentage was calculated by dividing the number of cells with evidence of P-L fusion by the total number of cells of five images for each condition. Data in (D) and (C) represent the mean ± SEM of four subjects, each with duplicate wells. *p<0.05, **p<0.01, and ***p<0.001 with one-way ANOVA test. CS=cigarette smoke, *MTB*=*Mycobacterium tuberculosis*, NS=non-significant.

An alternative approach to assess bacterial burden was employed in which MDM alone, MDM + unexposed Tregs, and MDM + CS extract-exposed Tregs were infected with GFP-*MTB* H37Rv for 2 and 4 days and the percentage of GFP-*MTB*-infected MDM determined by fluorescent microscopy. Compared to MDM alone, co-culture of Tregs with MDM increased the percentage of MDM containing GFP-labeled *MTB*, which further increased with the addition of Tregs that were pre-incubated with CS extract (**Figure 1B**). CS extract itself did not directly enhance the growth of *MTB* in the absence of macrophages (**Figure 1C**). We conclude that CS extract-exposed Tregs increased the burden of *MTB* in macrophages.

### CS extract-exposed Tregs decreased P-L fusion in *MTB*-infected MDM

An immune evasive mechanism of *MTB* is the inhibition of phagosome-lysosome (P-L) fusion. To determine whether CS extract-exposed Tregs affect P-L fusion in *MTB*-infected macrophages, MDM were cultured alone or co-cultured with autologous Tregs that had been previously exposed to medium alone or with 5% CS extract for 18-24 hours; this was followed by infection with GFP-*MTB* H37Rv for 6 hours and quantitation of P-L fusion. Compared to GFP-*MTB*-infected MDM alone, co-incubation of MDM with Tregs reduced the co-localization of GFP-*MTB* with lysosomes, with even further decrease when MDM were co-cultured with CS extract-exposed Tregs (**Figure 1D/E**).

### CS extract-exposed Tregs decreased autophagosome formation and maturation of *MTB*-infected MDM

To quantify autophagosome formation, MDM, MDM + unexposed Tregs, and MDM + CS extract-exposed Tregs infected with *MTB* for 18 hours were immunostained with anti-LC3-II antibody and Cy3-tagged goat anti-rabbit antibody. MDM co-incubated with CS extract-exposed Tregs had significantly fewer LC3-II(+) puncta than MDM alone or MDM + unexposed Tregs (**Figure 2A/B**). These findings indicate that in the presence of CS extract-exposed Tregs, *MTB*-infected MDM had either decreased autophagosome formation, increased autophagosome-lysosome fusion with degradation of autophagosome content, or both.

**Figure 2.**
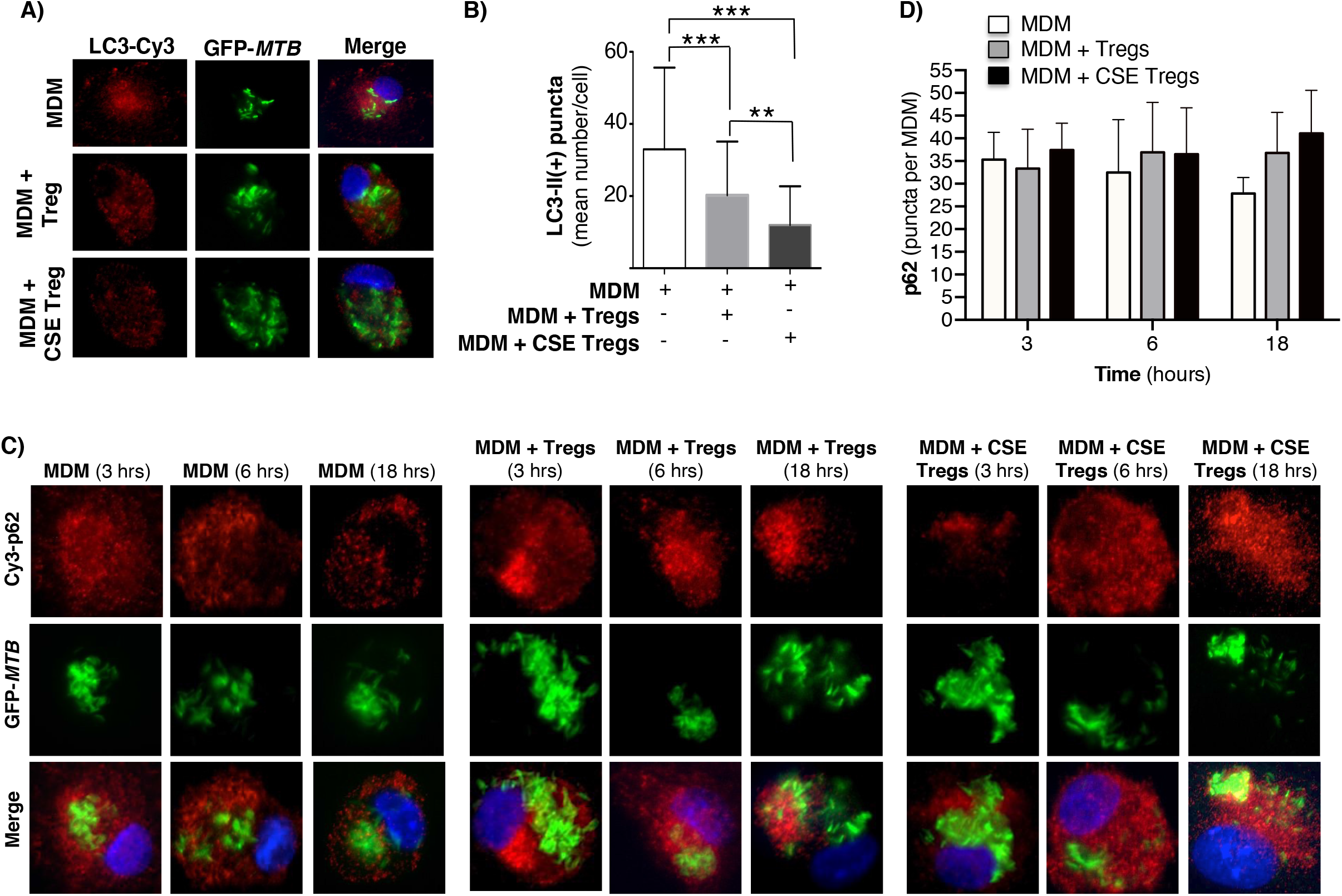
Cigarette smoke extract-exposed T regulatory cells decreases autophagosome formation. **(A)** Monocyte-derived macrophages (MDM) were incubated in medium alone or cocultured with unexposed or cigarette smoke (CS) extract-exposed T regulatory cells (Tregs), followed by infection with green fluorescent protein-labeled *Mycobacterium tuberculosis* (GFP-*MTB*) for 18 hours and assayed for LC3-II expression by immunofluorescence. **(B)** Mean LC3-II positive puncta were calculated by counting the individual puncta of *MTB*-infected MDM (150 cells per condition). **(C)** MDM were incubated in medium alone or co-cultured with unexposed or 5% CS extract-exposed Tregs, followed by infection with GFP-*MTB* for 3, 6 and 18 hours, and assayed for p62 expression by immunofluorescence. Data represent the mean ± SEM of four independent experiments. **(D)** Mean p62(+) puncta were calculated by counting the individual puncta of *MTB*-infected MDM (50 cells per condition). Data represent the mean ± SEM of three independent experiments, each with duplicate wells. **p<0.01 and ***p<0.001 with one-way ANOVA test.

To analyze which of these two autophagic processes is more dominant in MDM + CS extract-exposed Tregs, expression of the cargo protein sequestosome-1 (p62) – located within autophagosomes and degraded upon fusion with lysosomes – was temporally quantified. MDM, MDM + unexposed Tregs, and MDM + CS extract-exposed Tregs infected with *MTB* for 3, 6, and 18 hours were immunostained for p62. With MDM alone, there was a modest decrease in p62(+) autophagosomes with increasing time of infection, suggesting increased autophagosome-lysosome fusion (**Figure 2C/D**, open bars). However, in the presence of either unexposed Tregs or CS extract-exposed Tregs, the number of p62(+) autophagosomes did not decrease but there was a modest increase, suggesting a component of blockade in autophagosome-lysosome fusion in the presence of Tregs. Examination of the 18-hour time point shows a modest stepwise increase in p62(+) autophagosomes with MDM, MDM + unexposed Tregs, and MDM + CS extract-exposed Tregs (**Figure 2D**, last three bars), indicating decreased autophagosome-lysosome fusion in the presence of Tregs, especially those that were exposed to CS extract.

To further validate these findings, we quantified p62(+) puncta in the MDM, MDM + unexposed Tregs, and MDM + CS extract-exposed Tregs at 18 hours after infection with *MTB* in the presence of rapamycin, a known inducer of autophagy (27). As previously shown, there was a successive increase in p62 puncta with MDM alone, MDM + unexposed Tregs, and MDM + CS extract-exposed Tregs (**Figure 3B**, open bars). In contrast, in MDM plus either unexposed or CS extract-exposed Tregs treated with rapamycin, there was no increase in p62(+) puncta compared to cells not treated with rapamycin (**Figure 3B**, compare gray bars vs. respective open bars). Because rapamycin did not increase the number of autophagosomes in co-culture of MDM + CS extract-exposed Tregs, it suggests CS extract-exposed Tregs are inhibiting the signaling pathway that leads to autophagosome formation at a site distal to where rapamycin is augmenting autophagosome formation (**Figure 3C**).

**Figure 3.**
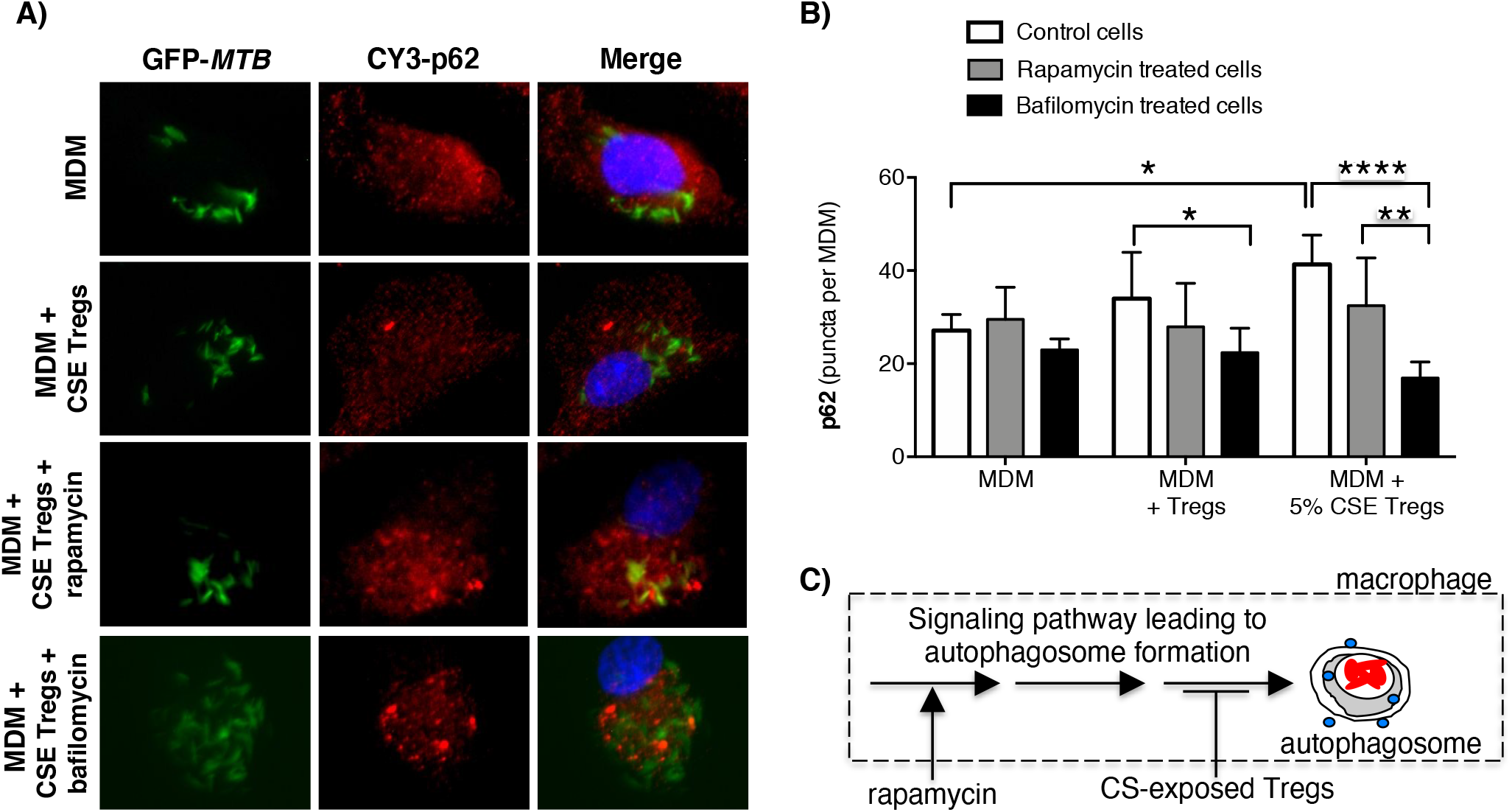
Cigarette smoke extract-exposed T regulatory cells decreases autophagosome formation at a site distal to rapamycin induction of autophagosomes. **(A)** Monocyte-derived macrophages (MDM) were incubated in medium alone or co-cultured with unexposed Tregs or cigarette smoke (CS) extract-exposed Tregs ± rapamycin or bafilomycin [100 nM] for 18 hours, followed by quantifying p62 expression by immunofluorescence**. (B)** Mean p62 puncta for a total of 50 cells per condition for the indicated conditions. **(C)** Based on the results, we hypothesize that CS extract inhibits autophagy at a site distal to where rapamycin induces autophagosome formation. Data represent the mean ± SEM of three independent experiments, each with duplicate wells. *p<0.05, **p<0.01, and ****p<0.0001 with one-way ANOVA test.

To determine whether CS extract-exposed Tregs are inducing autophagosome-lysosome fusion (= autophagosome maturation) to account for the decreased LC3-II(+) autophagosomes, we performed the aforementioned p62 assay in the absence or presence of bafilomycin, which inhibits both lysosome acidification and autophagosome-lysosome fusion (28). We hypothesized that if CS extract-exposed Tregs were inducing autophagosome-lysosome fusion, then in the presence of bafilomycin, we would expect to see a relative increase in the number of p62(+) puncta because bafilomycin would be inhibiting the degradation of p62(+) autophagosomes. Unexpectedly, addition of bafilomycin decreased the number of p62(+) puncta, especially in MDM + 5% CS extract-exposed Tregs (**Figure 3A/B**, last open vs. black bars). However, the size of the p62(+) puncta with bafilomycin treatment were noticeably larger than seen in MDM without bafilomycin treatment, suggesting autophagosome aggregation (**Figure 3A and Supp Figure 2A**). Using FIJI, the p62(+) puncta were measured by area and fluorescence using a Corrected Total Cell Fluorescence equation provided by Luke Hammond from The University of Queensland, Australia (29). These analyses showed that the total area fluorescence of the p62(+) puncta were not significantly different with or without bafilomycin (**Supp Figure 2B/C**). These findings support the paradigm that co-incubation of MDM with CS-exposed Tregs reduces autophagosome number by inhibiting their formation (as noted by decreased LC3-II autophagosomes) but also a component of decreased autophagosome maturation (as noted by increased p62(+) marker to the autophagosomes).

### Cytokine expression in *MTB*-infected MDM ± unexposed and CS extract-exposed Tregs

The supernatant of the MDM, MDM + unexposed Tregs, and MDM + CS extract-exposed Tregs used to obtain the aforementioned *MTB* CFU were saved and assayed for TNF, IL-10, and TGFβ. Compared to MDM alone, co-culture with unexposed Tregs modestly decreased TNF and increased IL-10 and TGFβ at 2 and 4 days after culture (**Figure 4A-C**). Co-culture of MDM with Tregs exposed to CS extract further decreased TNF level and increased IL-10 and TGFβ (**Figure 4A-C**).

**Figure 4.**
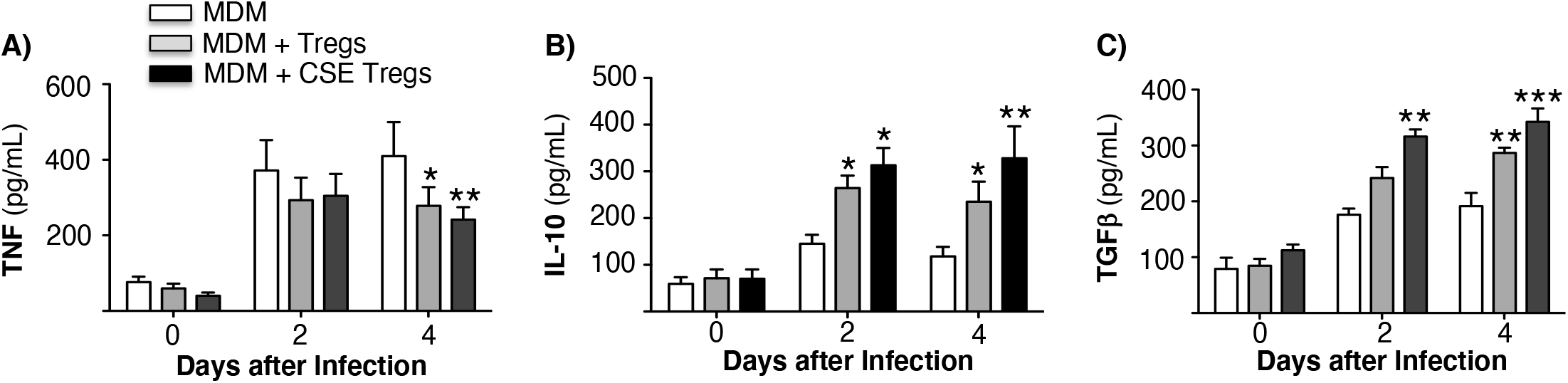
Cytokine expression by *Mycobacterium tuberculosis*-infected macrophages ± unexposed or cigarette smoke extract-exposed T regulatory cells. Monocyte-derived macrophages (MDM) and T regulatory cells (Tregs) were prepared from four different donors. Unexposed or cigarette smoke (CS) extract-exposed Tregs were added to MDM infected with *MTB*. These co-cultures and MDM alone infected with *MTB* were cultured for 1 hour, 2 and 4 days and the supernatants were obtained and measured for **(A)** TNF, **(B)** IL-10, and **(C)** TGFβ. Data shown are mean ± SEM of four independent experiments with each performed in duplicate wells. *p<0.05, **p<0.01, ***p<0.001. *MTB*=*Mycobacterium tuberculosis*

### CS extract induction of CTLA-4 on Tregs impairs MDM control of *MTB* infection

In human lungs, CS exposure promotes Treg function (20, 22, 23). A plausible mechanism by which this occurs is the ability of CS to induce PD-L1/2 on antigen presenting cells, resulting in increased engagement to PD-1 on Tregs (30, 31); *i.e*., whereas engagement of PD-1 and CTLA-4 on T cells to PD-L1/2 and B7 on macrophages, respectively, inhibits the activity of Foxp3-negative T effector cells, these same molecular interactions enhance the suppressive activity of Tregs (5, 32). Thus, to begin to determine the mechanism by which CS extract affects the ability of Tregs to impair macrophage control of *MTB* infection, we exposed Tregs to CS extract for 18 hours and quantified PD-1 and CTLA-4 expression. As shown in **Figure 5A**, CS extract increased the expression of CTLA-4 but not PD-1 in primary human Tregs.

**Figure 5.**
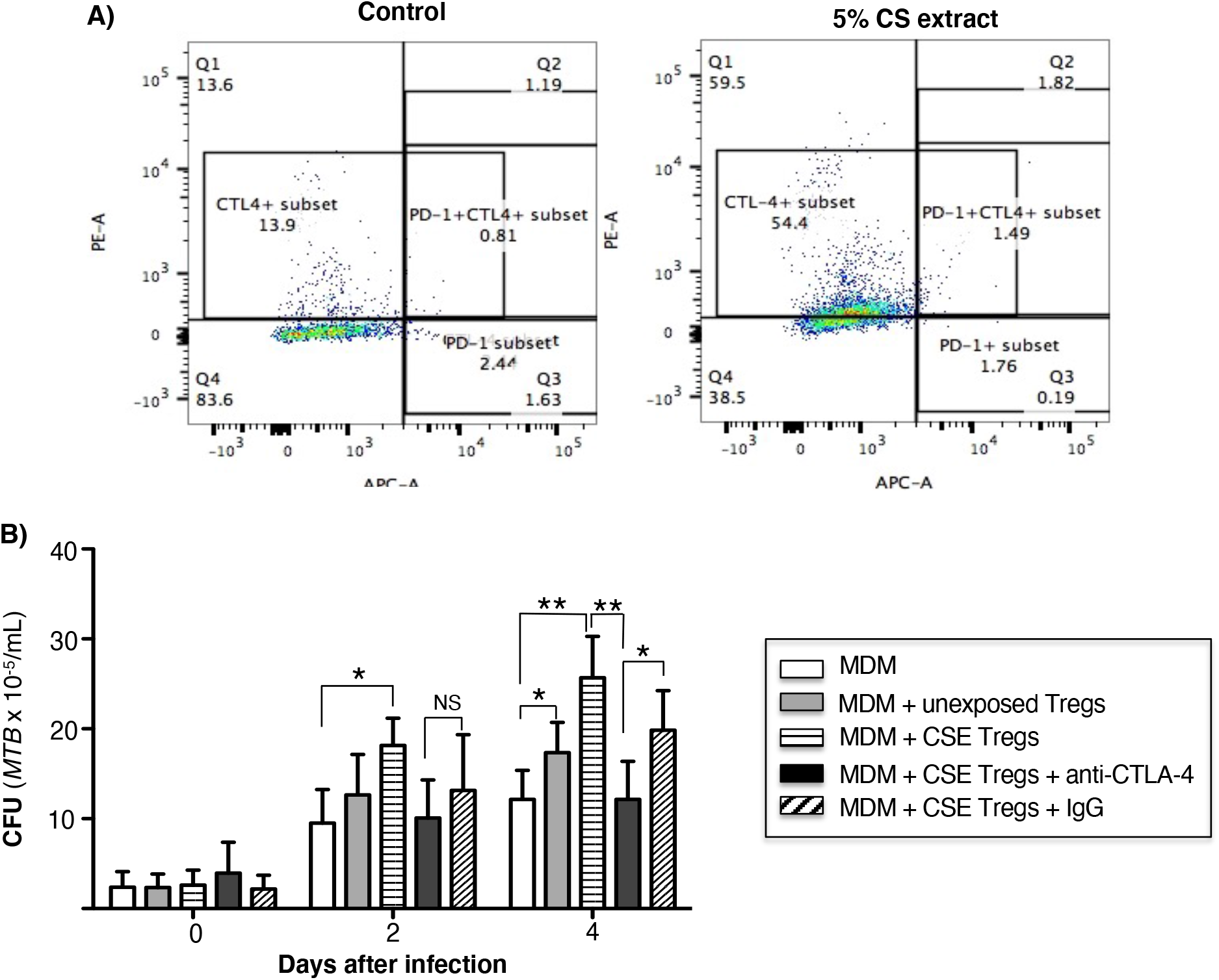
Cigarette smoke extract induction of CTLA-4 in T regulatory cells increases their ability to impair macrophage control of *Mycobacterium tuberculosis*. **(A)** Unexposed or cigarette smoke (CS) extract-exposed T regulatory cells (Tregs) of a healthy subject were stained with PE-conjugated anti-CTLA-4 and APC-conjugated anti-CD279 (PD-1) and flow cytometry was performed. **(B)** Monocyte-derived macrophages infected with *MTB* were cultured alone or co-cultured with unexposed or CS extract-exposed Tregs ± anti-CTLA-4 or non-immune IgG for 1 hour, 2, and 4 days. After the indicated times, intracellular *MTB* was quantified. Data shown are the mean ± SEM of cells derived from four different donors and performed in duplicates. *p<0.05, **p<0.01, *MTB=Mycobacterium tuberculosis*, NS=non-significant.

To determine the role that CS extract-induced CTLA-4 on Tregs may play in impairing MDM control of *MTB*, we incubated CS extract-exposed Tregs with 5 μg/mL non-immune IgG or with 5 μg/mL anti-CTLA-4 antibody before combining the Tregs with *MTB*-infected MDM (33). As shown in **Figure 5B**, macrophages co-cultured with CS extract-exposed Tregs have greater *MTB* burden than MDM alone or MDM + unexposed Tregs. However, in the presence of anti-CTLA-4 antibody, there was a partial but significant abrogation of the increase in *MTB* burden in co-culture of MDM + CS extract-exposed Tregs; such abrogation was not seen with non-immune IgG. These findings indicate that CS extract induction of CTLA-4 on Tregs is a mechanism by which CS extract-exposed Tregs impair macrophage control of *MTB* infection.

### Adoptive transfer of Tregs from CS-exposed mice increased *MTB* burden in recipient mice

Following adoptive transfer of Tregs from air-exposed and CS-exposed B6.PL(Thy1.1) donor mice into Treg depleted Foxp3^+^GFP^+^DTR^+^ (Thy1.2) mice, the latter recipient mice were infected with HN878-W-Beijing *MTB* for 1, 30, and 60 days. Due to the immunocompetent phenotype of these mouse strains, a hypervirulent strain of *MTB* was chosen for infection to generate a productive infection. As an additional control, Foxp3^+^GFP^+^DTR^+^ (Thy1.2) mice, in which no Treg depletion or transfer was performed, were also infected with HN878 *MTB*. The lungs and spleen were harvested, processed, and the CFU of the organs were determined at the indicated times. Compared to control mice and mice that received Tregs from air-exposed mice, those that received Tregs from CS-exposed mice had significantly greater CFU in both the lungs and spleens but only at Day 60 after infection (**Figure 6A-B**).

**Figure 6.**
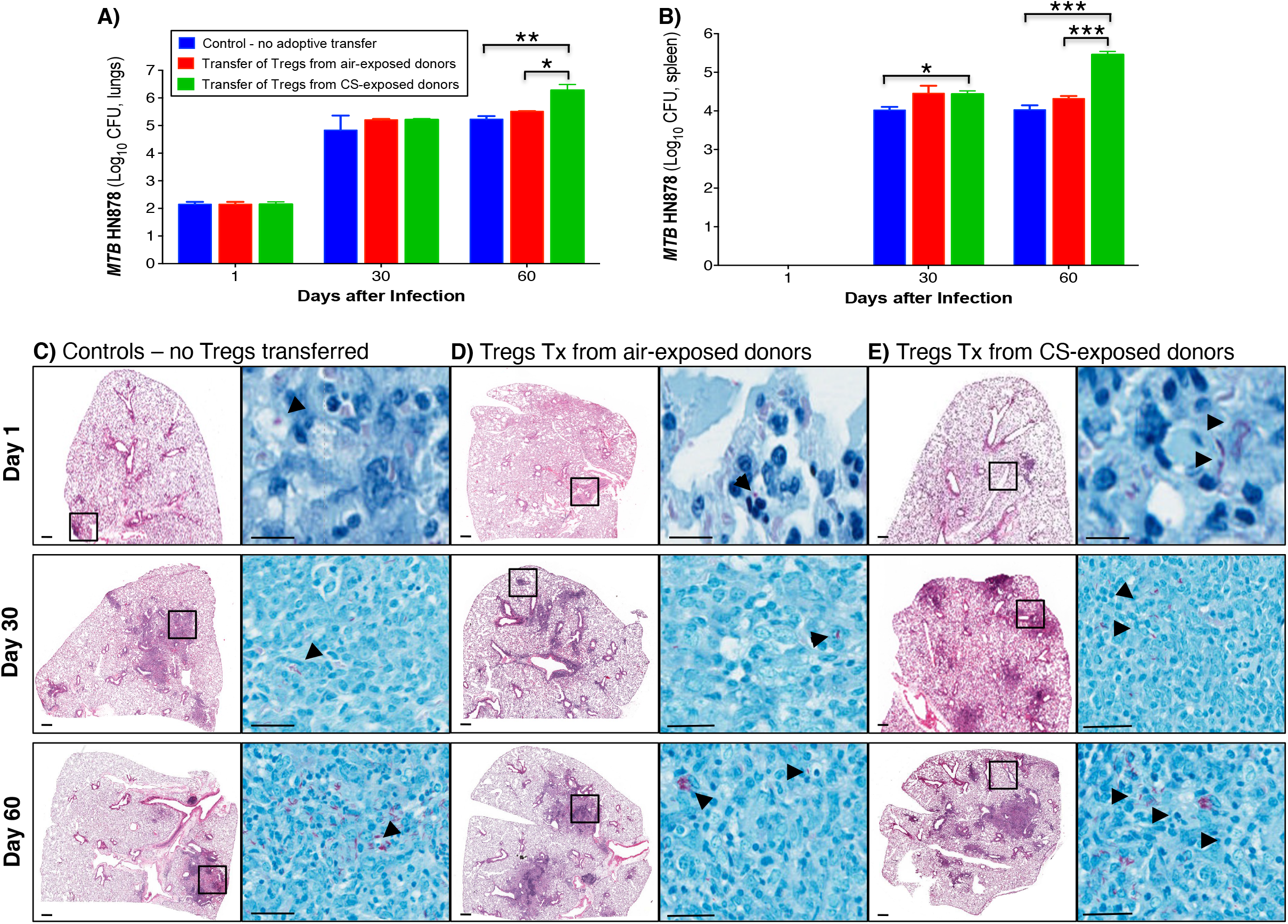
Adoptive transfer of T regulatory cells from cigarette smoke-exposed mice have greater burden of *Mycobacterium tuberculosis* in the lungs and spleen. **(A)** T regulatory cells (Tregs) were isolated from two groups of Thy1.1 mice; one group was exposed to cigarette smoke (CS), while the other was exposed to ambient air. The isolated Tregs from unexposed and CS-exposed Thy1.1 mice were adoptively transferred into recipient, Treg-depleted Thy1.2 mice. A third group of control mice did not undergo Treg depletion or adoptive transfer. All three groups of mice were then infected with *MTB* HN878. The mice from each group were sacrificed at the one, 30 and 60 days after infection. The lungs of mice were homogenized to quantify CFU. **(B)** The spleens of mice were also homogenized to determine CFU. Representative histopathologic photomicrograph (H&E and acid-fast stains) are shown of **(C)** control mice that were infected with *MTB* alone and recipient mice that had previously received Tregs from **(D)** air- or **(E)** CS-exposed mice and then infected with *MTB*. Data shown are the mean ± SEM of cells derived from five mice. *p<0.05, **p<0.01, ***p<0.001. *MTB*=*Mycobacterium tuberculosis*.

### Lung histopathology of mice that received Treg from CS-exposed mice revealed larger, less-discrete lung lesions

Histopathology of control mice without Treg transfer, and mice with transferred Tregs from air- or CS-exposed mice was evaluated 1, 30 and 60 days after HN878 *MTB* infection. Compared to control mice and mice recipient of Tregs from air-exposed donors, mice that received Tregs from CS-exposed donors demonstrated increased size and number of granuloma-like lesions with increased granulocyte infiltration, and presence of acid-fast bacilli (**Figure 6C-E, arrows**).

### Immunophenotyping of murine lung macrophages and dendritic cells

Lung macrophages and dendritic cells (DC) of uninfected mice, *MTB*-infected mice, and Foxp3^+^GFP^+^DTR^+^ (Thy1.2) mice recipient of air- or CS-exposed Tregs followed by *MTB* infection were immunophenotyped for intracellular TNF, IL-12, and IL-10 at 1, 30 and 60 days after *MTB* infection. Compared to no infection, *MTB* infection increased the number of TNF^+^ lung macrophages and DC at 30 and 60 days after infection (**Figure 7A/B**). At 60 days after infection, there was decreased number of TNF^+^ macrophages in mice transferred Tregs from CS-exposed mice although this did not reach statistical significance (**Figure 7A**). At 60 days post infection, there was also a trend toward decreased TNF^+^ DC in mice transferred Tregs from both air- or CS-exposed mice (**Figure 7B**).

**Figure 7.**
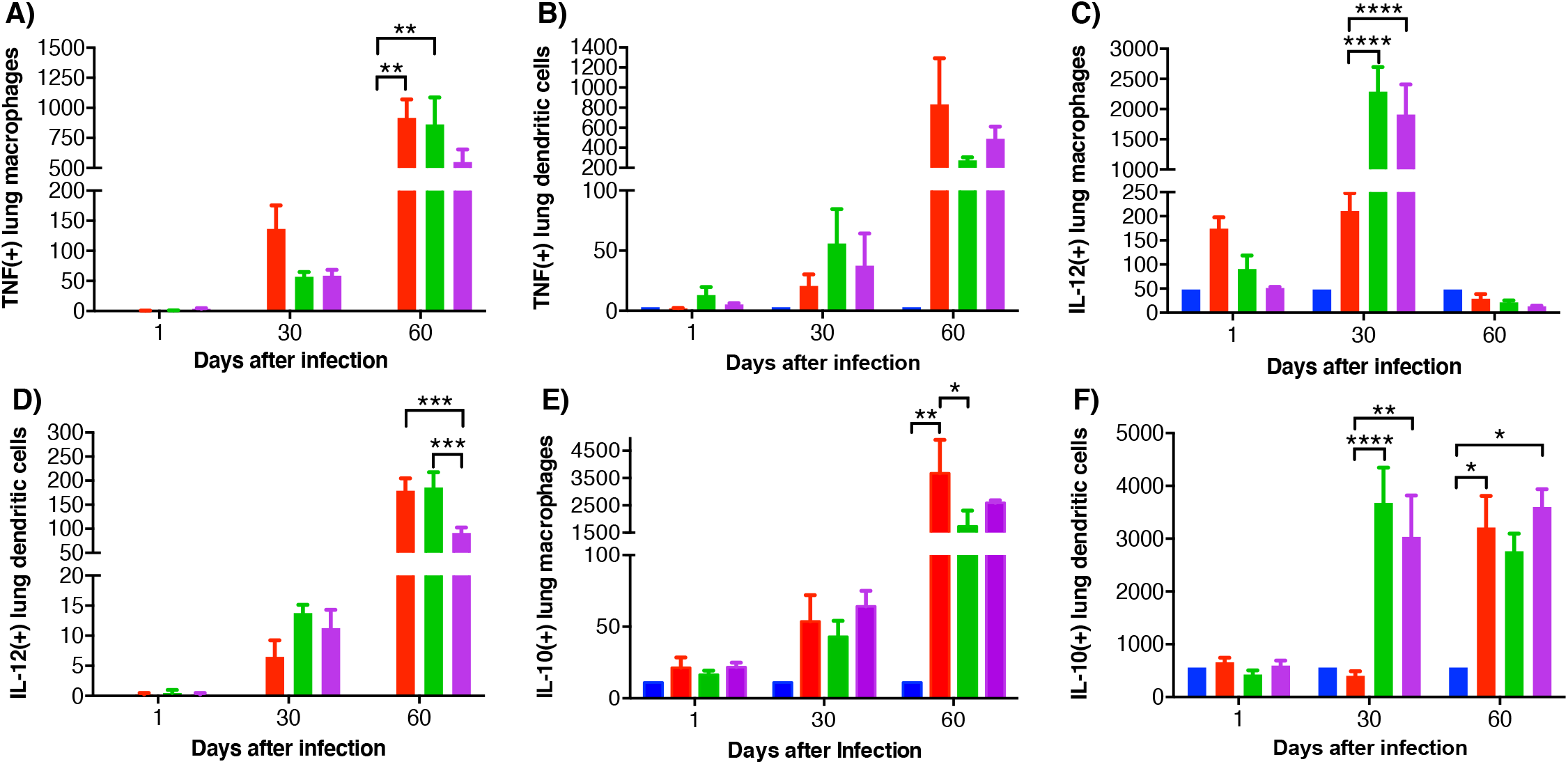
Intracellular cytokine analysis of murine lung macrophages and DCs. Lung macrophages and dendritic cells from uninfected mice, *MTB*-infected mice, and *MTB*-infected mice with adoptive transfer of Tregs from either air-exposed or CS-exposed mice were stained for **(A/B)** TNF, **(C/D)** IL-12, and **(E/F)** IL-10. Data shown are the mean ± SEM of cells derived from five mice. *p<0.05, **p<0.01, ***p<0.001, ****p<0.0001. *MTB=Mycobacterium tuberculosis*.

Compared to uninfected mice, *MTB* infection alone increased the number of IL-12^+^ lung macrophages, which further increased with the transfer of either air- or CS-exposed Tregs at 30 days post-infection (**Figure 7C**). Interestingly, at 60 days post-infection, IL-12^+^ lung macrophages decreased to levels even lower than at 1 day after infection. In contrast, IL-12^+^ DC progressively increased temporally with all three *MTB*-infected conditions (**Figure 7D**); in the mice that received Tregs from CS-exposed mice, there was a significantly lower number of IL-12^+^ DC compared to mice that were uninfected, *MTB* infection only, and mice that received Treg from air-exposed mice and *MTB* infected (**Figure 7D**).

In analyzing lung macrophages and DC that expressed IL-10, there was a temporal increase of IL-10^+^ lung macrophages and DC with *MTB* infection (except for DC at Day 30 after infection) (**Figure 7E/F**). Mice that received Tregs from either air- or CS-exposed mice have decreased IL-10^+^ macrophages at Day 60 compared to *MTB*-infected mice alone without Treg transfer (**Figure 7E**). In contrast, mice that received either air- or CS-exposed Tregs and *MTB*-infected have marked infiltration with IL-10^+^ DC at Day 30 compared to mice infected with *MTB* only, but this difference was lost at Day 60 (**Figure 7F**).

While CTLA-4 is canonically expressed on T cells, it is also present on human monocytes and mature monocyte-derived DC following stimulation with lipopolysaccharide, poly:IC, and various cytokines; CTLA-4 on DC also function to inhibit CD4^+^ T cell proliferation *via* CTLA-4-mediated production of IL-10 by the DC (34). Thus, we quantified CTLA-4 expression in the mouse lungs and found that with *MTB* infection, there was a temporal increase in CTLA-4^+^ macrophages with *MTB* infection with or without Treg transfer (**Figure 8A**). However, in the mice that received air-exposed Tregs, there was a lower amount of CTLA-4^+^ macrophages at Day 60, but in the mice that received Tregs from CS-exposed mice, there was higher number of CTLA-4^+^ macrophages (**Figure 8A**). There was also a temporal increase of CTLA-4^+^ DC with all infections – except for Day 30 with the only *MTB*-infected mice (**Figure 8B**). In contrast to that seen with lung macrophages, there was a significant decrease of CTLA-4^+^ DC in the mice that received Tregs from CS-exposed mice at Day 60 compared to mice with *MTB* infection alone. We conclude that in mice that received Tregs from CS-exposed mice, there was greater decrease in the number of macrophages and DC that expressed TNF and IL-12 and the number of CTLA-4^+^ DC.

**Figure 8.**
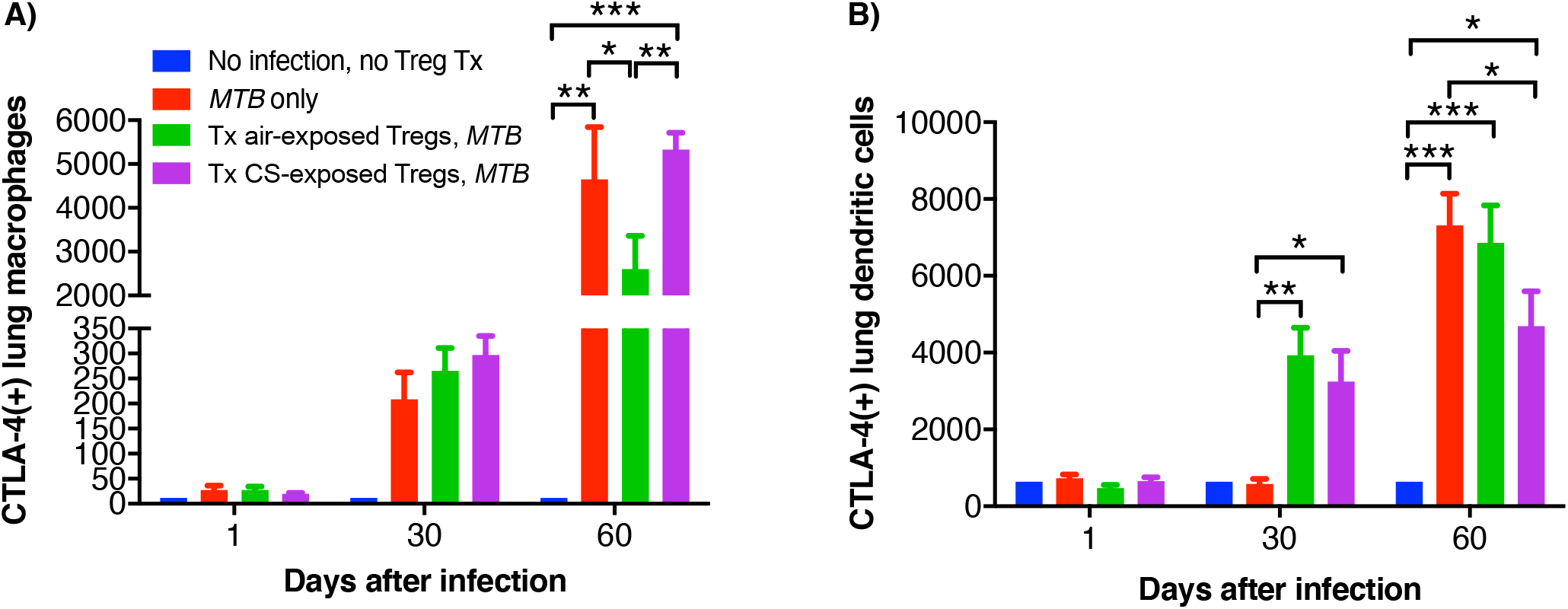
CTLA-4 expression in the lung macrophages and DC of the recipient mice. At the indicated times, uninfected mice, after *MTB*-infected Thy1.2 mice, and mice that received T regulatory cells (Tregs) from either air- or CS-exposed Thy1.1 mice and subsequently infected with *MTB* were sacrificed and CTLA-4^+^ lung **(A)** macrophages and **(B)** dendritic cells were quantified by flow cytometry. Data shown are the mean ± SEM of cells derived from five mice. *p<0.05, **p<0.01, ***p<0.001. *MTB=Mycobacterium tuberculosis*.

### Immunotyping of murine lung CD4^+^ T cells

The general gating strategy for T cells is as shown in **Figure 9A**. Examining the phenotypic subsets of T cells in the lungs revealed a temporal increase in CD4^+^IFN*γ*^+^ T cells in the lungs of all the three mouse groups infected with *MTB* compared to uninfected mice (**Figure 9B**). But at Day 60, the increase of CD4^+^IFN*γ*^+^ T cells in mice that received Tregs from CS-exposed mice and infected with *MTB* was less than mice with *MTB* infection alone or that received Tregs from air-exposed mice + *MTB* infection but this difference was not statistically significant.

**Figure 9.**
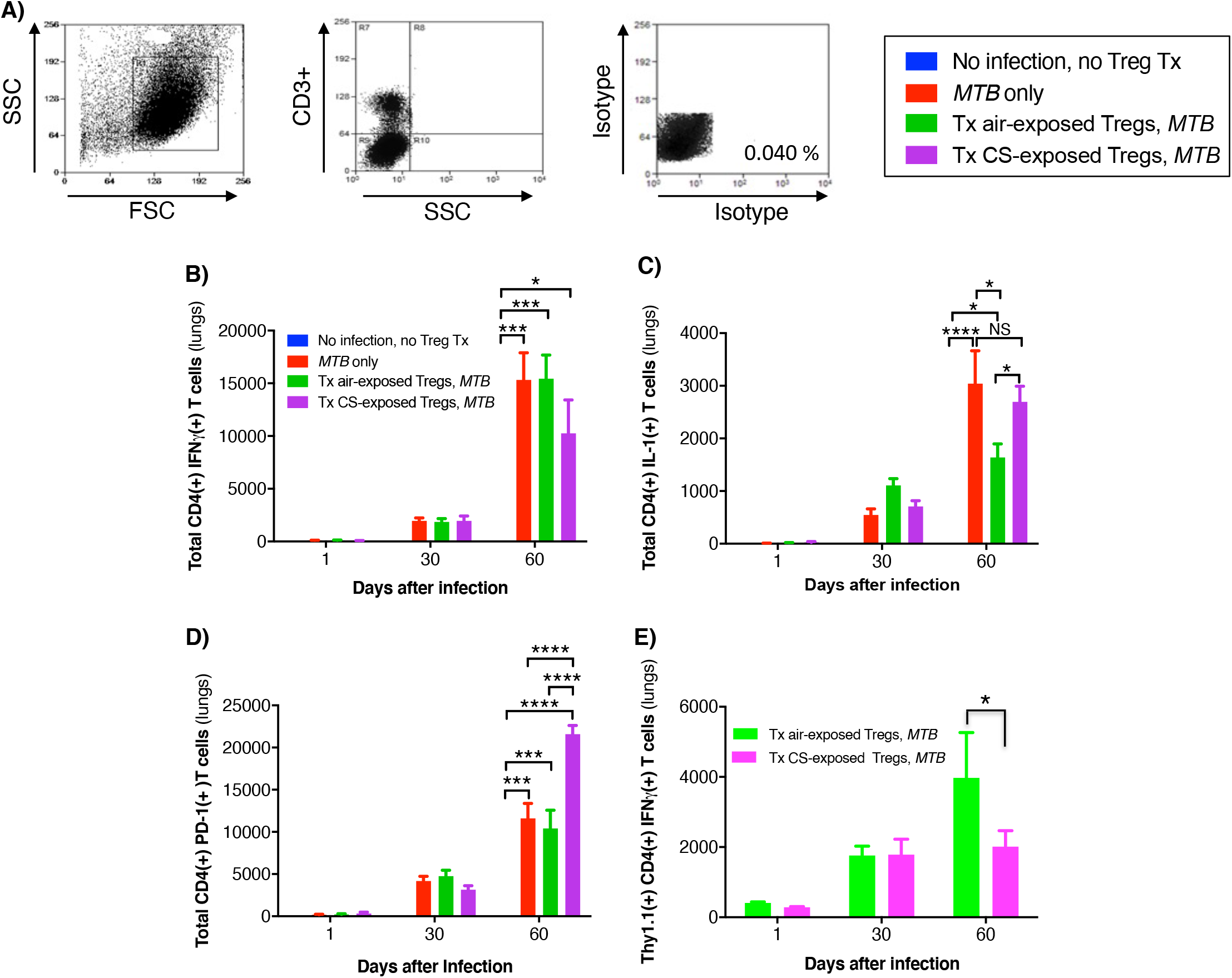
Cytokine expression of T cells in the lungs of mice that received adoptive transfer of T regulatory cells. **(A)** General gating strategy for T cells in the murine lungs. **(B)** CD4^+^IFN*γ*^+^ T cells, **(C)** CD4^+^IL-1^+^ T cells, and **(D)** CD4^+^PD-1^+^ T cells were quantified in the lungs of the uninfected mice, *MTB*-infected mice, and Thy1.2 mice that received T regulatory cells (Tregs) from air- or cigarette smoke (CS)-exposed Thy1.1 mice and then infected with *MTB*. **(E)** Thy1.1 CD4^+^IFN*γ*^+^ transferred Tregs quantified in the lungs of the recipient Thy1.2 mice at the indicated times. Data shown are the mean ± SEM of cells derived from five mice. *p<0.05, ***p<0.001, and ***p<0.0001. *MTB=Mycobacterium tuberculosis*.

Compared to uninfected mice, the number of CD4^+^IL-1^+^ T cells increased temporally in control mice and mice that received Tregs from air- or CS-exposed donor mice. The number of CD4^+^IL-1^+^ T cells was similar at Days 1 and 30 between all three groups of mice infected with *MTB* (**Figure 9C**). But by Day 60, mice that received air-exposed Tregs had the least number of IL-1^+^CD4^+^ T cells (**Figure 9C**). In examining an exhaustion marker of CD4^+^ T cells, mice that received Tregs from CS-exposed mice had significantly more PD-1-positive T cells at Day 60 than either mice that were infected with *MTB* only or that received Tregs from air-exposed mice and were subsequently infected with *MTB* (**Figure 9D**).

Quantifying the total number of CD4^+^ Tregs in the four groups of mice (uninfected, *MTB*-infected alone, and *MTB*-infected mice recipient of Tregs from air- or CS-exposed mice), we found that compared to uninfected mice, *MTB*-infected mice had significantly more total Tregs, which peaked at Day 30 after infection (**Supp Figure 3A**). While there was a lower number of total CD4^+^ Tregs at Day 30 after infection in the mice that received Tregs from CS-exposed mice, this was not statistically significant. Similarly, there was a trend of increased number of CD4^+^IL-10^+^ Tregs at Day 60 after *MTB* infection in the mice that received Tregs from either air- or CS-exposed mice (**Supp Figure 3B**). Since there is a small population of Tregs that can produce IFN*γ* (35–37), we analyzed the population of transferred Thy1.1^+^CD4^+^IFN*γ*^+^ Tregs from the air- or CS-exposed Thy1.1 mice in the lungs of the recipient Thy1.2 mice. While there was no difference in the population of these Tregs in the lungs at Day 1 or 30, by Day 60 the recipient Thy1.2 mice had fewer donor Thy1.1^+^IFN*γ*^+^ Tregs from the CS-exposed mice compared to that from the air-exposed mice (**Figure 9E**). Furthermore, the temporal increase in the donor CD4^+^IFN*γ*^+^ Tregs indicates replication of these donor Tregs in the recipient mice over the course of the infection as has been previously described (38).

### Immunophenotyping of murine splenic macrophages, dendritic cells, and T cells

Splenic macrophages and DC were also analyzed 2, 30, and 60 days after *MTB* infection in control mice infected with *MTB* only and in mice that had received Tregs from air- or CS-exposed mice before *MTB* infection. Uninfected mice also served as an additional control. Compared to uninfected mice, the three mouse groups infected with *MTB* demonstrated an increase in intracellular TNF^+^, IL-12^+^, and IL-10^+^ macrophages (**Supp Figures 4A-C**) and DC (**Supp Figure 4D-F**). With the time points examined, the number of such macrophage and DC phenotypes were highest at 60 days after *MTB* infection except for the IL-12^+^ macrophages, which peaked at Day 30 (**Supp Figure 4B**). Similarly, the three murine groups infected with *MTB* had similar numbers of CTLA-4^+^ macrophages (**Supp Figure 4G**) and dendritic cells (**Supp Figure 4H**), with both cell type numbers highest at 60 days after infection.

Analysis of the total number of splenic CD4^+^ T cell subsets revealed that, compared to uninfected mice, the number of T cells significantly increased at 30 and 60 days in the three mouse groups that were infected with *MTB* (**Supp Figure 5**). More specifically, the total number of IFN*γ*^+^ T cells increased progressively from Day 2 to Day 60 after infection in control mice that were only infected with *MTB* and in *MTB*-infected mice that received Tregs from air-exposed mice. In contrast, the temporal increase in the number of IFN*γ*^+^ T cells was attenuated in the animals that received Tregs from CS-exposed mice (**Supp Figure 5A**). Similarly, compared to uninfected mice, there was an increase in the number of IL-1^+^ T cells in the three mouse groups infected with *MTB* (**Supp Figure 5B**). The number of such IL-1^+^ T cells peaked at Day 30 in the mice infected with *MTB* alone and mice that received Tregs from air-exposed mice whereas they peaked at Day 60 in the mice that received Tregs from CS-exposed mice and subsequently infected with *MTB* (**Supp Figure 5B**). In examining the CD4^+^ T cell subset expressing the PD-1 exhaustion marker, there was a progressive temporal increase in all three groups of mice infected with *MTB* (**Supp Figure 5C**). However, there was significantly greater number of PD-1^+^ splenic T cells in the *MTB*-infected mice that received Tregs from CS-exposed mice (**Suppl Figure 5C**).

In analyzing the splenic Tregs, there was an increased in total number at Days 30 and 60 in the three mouse groups infected with *MTB* compared to uninfected mice (**Supp Figure 5D**). Compared to *MTB* infection alone, those that received air-exposed mice had a robust rise in Tregs at Day 30, which decreased to near basal levels at Day 60. In contrast, there was a progressive increase in Tregs in mice that received Tregs from CS-exposed mice. The number of IL-10^+^ Tregs temporally quantified in the spleen followed a very similar pattern to that observed with the total number of Tregs (**Supp Figure 5E**).

A summary of the effects of the experimental mouse conditions are listed in Table 1.

**Table 1.** Summary of the lung immune cell phenotype of the adoptive transfer murine studies of adoptive transfer of Tregs from CS-exposed mice compared to that of air-exposed mice.

| Significant difference | Trend | No effect |
| --- | --- | --- |
| • Decrease in IL-12+ dendritic cells (D60)<br>• Increase in CTLA4+ macrophages | • Decrease in CTLA4+ dendritic cells<br>• Decrease in IL-12+ macrophages<br>• Increase in IL-10+ macrophages | • IL-10+ dendritic cells<br>• TNFa+ macrophages and dendritic cells |
| • Increase in PD-1+ CD4+ T cells (D60)<br>• Increase in IL-1+ CD4+ T cells (D60) | • Decrease in IFNg+ CD4+ T cells<br>• Decrease in total IFNg+ T cells | • IL-10+ CD4+ Tregs |

**Table 2.** Summary of the splenic immune cell phenotype of the adoptive transfer murine studies of adoptive transfer of Tregs from CS-exposed mice compared to that of air-exposed mice.

| Significant difference | Trend | No effect |
| --- | --- | --- |
| • Increase in IL-1 Tcells (D60)<br>• Increase in total PD1 Tcells (D60) | • Increase in Tregs (D60)<br>• Increase of IL10 Tregs<br>• Decrease in IFNy Tcells<br>• Increase in PD1+ Tcells D60<br>•<br>• | • |
| • | • | • |

Decreased IL-12(+) dendritic cells

## DISCUSSION

While there is robust epidemiologic evidence that CS exposure is significantly associated with the development of active TB and with greater severity (2), less is known of the mechanisms by which this link occurs. CS has been shown to be polarized toward the M2/deactivated phenotype and to impair macrophage control of *MTB* (7, 9, 10, 39). Previous studies of mice exposed to CS and subsequently infected with *MTB* demonstrated similarly decreased influx of host-protective cells, particularly IFN*γ*-producing T_H_1 cells and M1 type macrophages, and an increased number of CD4^+^IL-4^+^ T cells (6, 8, 9). However, less is even known of how CS may affect the immunosuppressive cells such as Tregs in the context of TB.

Tregs are specialized CD4^+^ T cells that develop from naïve T cells in the presence of TFGβ. Tregs produce anti-inflammatory cytokines TGFβ, IL-35 and IL-10 (40–42). Through these cytokines, Tregs can suppress anti-TB immunity during the early stages of infection (14, 15, 17–19). Another immunosuppressive mechanism of Tregs is their ability to inhibit T effector cells, in part, through the secretion of granzyme B, which breach the cell walls of T effector cells, causing them to undergo apoptosis in a perforin-dependent fashion (43). Tregs may also suppress M1 macrophage activation and induce macrophage differentiation to the M2 phenotype (41, 44, 45). M2 macrophages exhibit impaired induction of pro-inflammatory cytokines and macrophage effector functions against *MTB* such as P-L fusion, autophagy, and apoptosis (46, 47). In addition, the cell surface protein LAG3 on Tregs binds to MHCII on DC, preventing further DC activation (48). The surface protein TIGIT found on T effector cells and Tregs also binds DC causing secretion of TGFβ and IL-10, immunosuppressive cytokines capable of inhibiting T effector cells and DC (49). However, Tregs may also be necessary at the later stages when *MTB* infection comes under control to prevent excessive inflammatory tissue damage even from CD4^+^IFN*γ*^+^ T cells (50, 51).

CS and nicotine are also known to increase the activation or production of Tregs (20–23). Using primary murine cells, we previously demonstrated that nicotine-exposed Tregs impaired murine macrophages from controlling an *ex vivo MTB* infection (11). This current study revealed that CS exposure of *only* primary human Tregs also impaired macrophage control of *MTB* infection through inhibiting macrophage effector functions against *MTB* including decreased phagosome-lysosome fusion and autophagosome formation and maturation.

While rapamycin is known to increase autophagosome formation by inhibiting mTOR, it could also indirectly drive autophagosome degradation – analogous to LeChatelier’s principle with chemical reaction by providing an increased number of the substrate (autophagosomes). Thus, if rapamycin-induced autophagosome degradation rate is greater than rapamycin-induced autophagosome formation rate and combined with decreased autophagosome formation due to the CS extract-exposed Tregs, one may not see an increase in autophagosomes with rapamycin and in fact may see a modest decrease, which is what was observed (Figure 3A/B). These pathways are made more complex by the finding that rapamycin can also increase p62 mRNA and protein expression in an autophagy independent pathway (27). We also found that in the presence of bafilomycin, an inhibitor of autophagosome-lysosome fusion, there was no significant difference in the total p62(+) puncta, which would be an expected finding if an increase in p62(+) puncta due to bafilomycin was countered by decreased autophagosome formation from the CS extract-exposed Tregs. We conclude from these autophagy experiments that CS extract-exposed Tregs reduction in macrophage autophagosome formation predominated over the decrease in autophagosome maturation.

One molecular mechanism for the increased Treg activity with CS or nicotine exposure is induction of CTLA-4 and PD-L1, the latter a ligand for PD-1 (30, 31, 52, 53). While engagements of CTLA-4 and PD-1 to their respective ligands deactivate effector T cells, they enhance Treg function, as supported by the observation that blockade of CTLA-4 and PD-1 are known to exacerbate autoimmune diseases through deactivation of Tregs (54, 55). But in human Tregs, we did not find an increase in the number of PD-1 positive Tregs but rather an increase in CTLA-4 positive Tregs with CS extract exposure. We further showed CS induction of CTLA-4 is a mechanism by which CS-exposed Tregs impair macrophage control of *MTB* infection.

The murine adoptive transfer studies corroborated the *in vitro* human cell data in that recipient Thy1.2 mice that received Tregs from donor CS-exposed mice had increased *MTB* burden in both the lungs and spleens compared to the two murine controls (*MTB* infection of naïve mice and of mice that received Tregs from air-exposed mice). Interestingly, the increase in CFU of the mice that received Tregs from CS-exposed mice was not observed until Day 60 in the lungs and spleens. While there was a significant increase in CFU with mice that received Tregs from CS-exposed mice in the spleen at Day 30, it was by ~one-half log of CFU increase.

The increased *MTB* burden in the animals that received Tregs from CS-exposed mice may be partly explained by the following immunophenotypic findings in these mice compared to the two control mice: significant decrease in IL-12^+^ DC, an increase in CTLA-4^+^ macrophages, and an increase in CD4^+^PD-1^+^ T cells and a trend toward increased IL-10^+^ macrophages and decreased TNF^+^ lung macrophages, IL-12^+^ macrophages, and total IFN*γ*^+^CD4^+^ T cells and Thy1.1^+^CD4^+^IFN*γ*^+^ transferred Tregs (**Table 1**). However, in recipient mice that received Tregs from donor CS-exposed mice, we found decreased number of CTLA-4-positive DC. CTLA-4 has been previously reported on macrophages (56, 57) and dendritic cells (58) and may play a role in inhibiting immune response. Thus, this finding seems paradoxical since a decrease in CTLA-4^+^ DC would not be expected to inhibit CD4^+^ T effector cell activation as much, which would relatively enhance host immunity against *MTB* (34).

The reduction in IFN*γ*^+^ T cells would be expected to reduce macrophage activation and MHCII expression. Unexpectedly, mice that received air-exposed Tregs had less CD4^+^IL-1^+^ T cells than those that received Tregs from CS-exposed mice since IL-1 is generally considered to be host-protective. Since a subset of Tregs can also produce IFN*γ* (35–37), it is interesting to note that there was a trend toward decreased IFN*γ*^+^ Tregs in the mice that received Tregs from CS-exposed donor mice than mice recipient of Tregs from air-exposed mice. Furthermore, in the animals that received Tregs from air-exposed mice, IFN*γ*^+^ Tregs increased temporally indicating multiplication of the transferred Tregs, which has been previously reported (38).

## SUMMARY

In conclusion, we found that isolated exposure of Tregs to CS (or CS extract) takes an active role in impairing human macrophage and mouse immunity against *MTB*. In primary human cells, CS extract-exposed Tregs impaired macrophage control of *MTB* by inhibiting phagosome-lysosome fusion and autophagosome formation and to a lesser extent by inhibiting autophagosome maturation. Mice that received Tregs from CS-exposed mice were also more compromised in controlling *MTB* infection due mainly to a decrease in effector innate and T cell phenotypes considered to be host-protective.

---

**Supplementary Figure 1.**
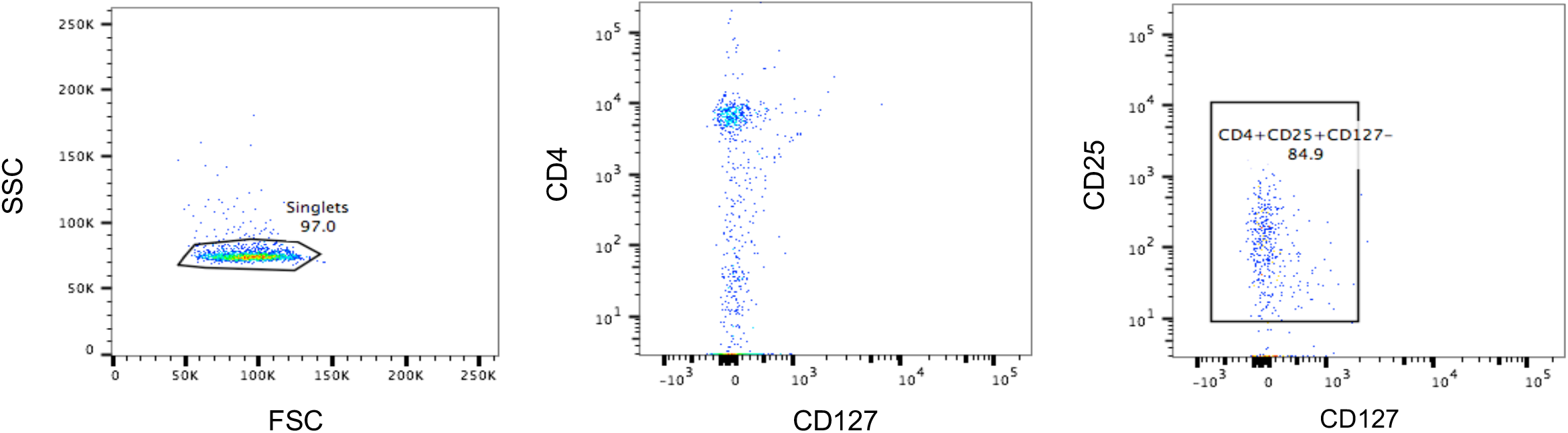
Human T regulatory cell isolation. T regulatory cells (Tregs) were isolated from peripheral blood mononuclear cells using Miltenyi Biotec’s Human Treg Isolation Kit II. Following isolation, the cells were stained for CD127-BB515, CD25-PE, and CD4-APC-CY7 to confirm Treg enrichment.

**Supplementary Figure 2.**
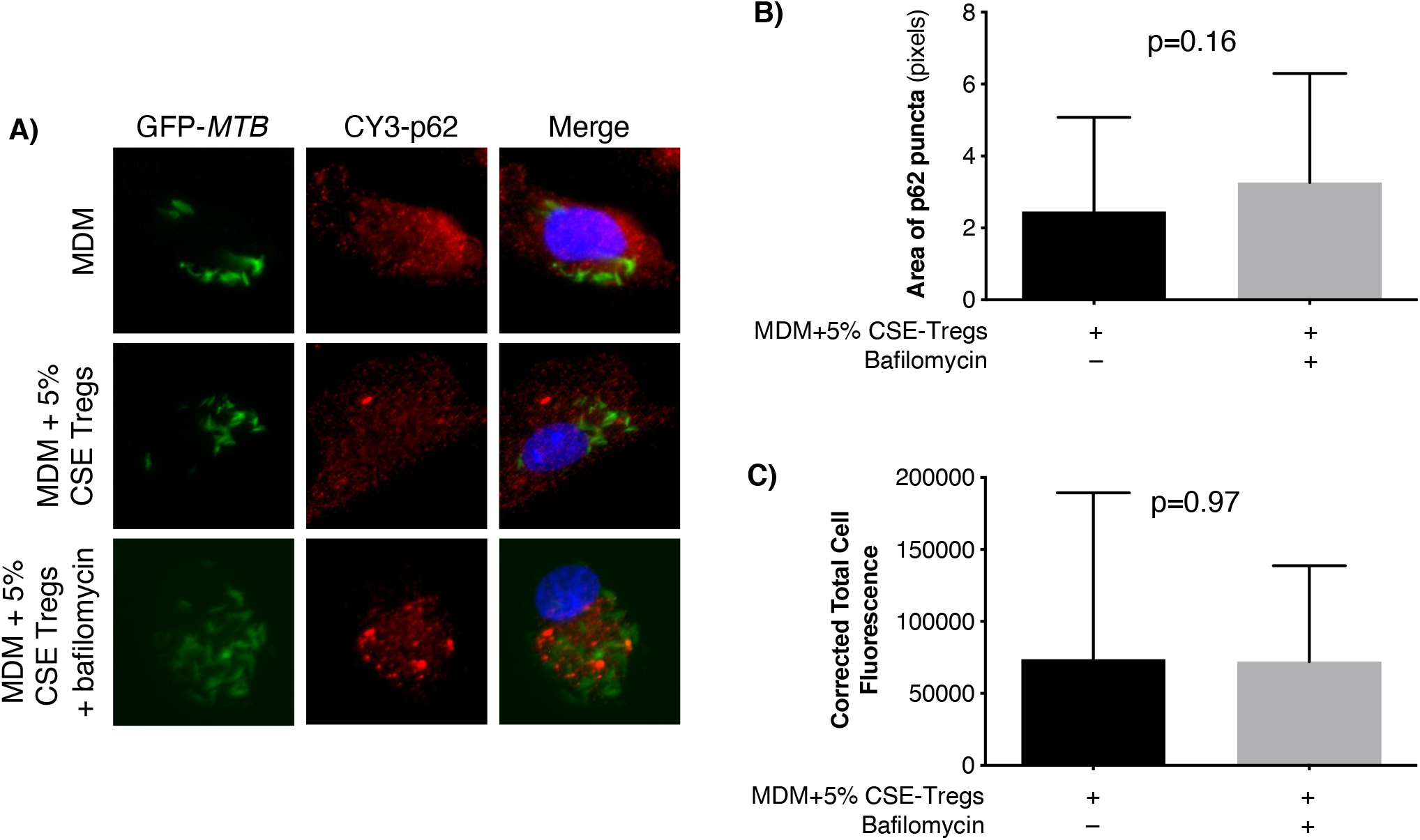
Puncta size and corrected total cell fluorescence of bafilomycin treated cells. **(A)** Human monocyte-derived macrophages were incubated as previously stated, followed by infection with a 10:1 MOI of GFP-*MTB* with or without bafilomycin for 18 hours and assayed for p62 expression immunofluorescence. **(B)** The area of p62 puncta was measured using FIJI software. A total of 300 puncta were measured, the graph depicts the mean area of the p62(+) puncta in MDM co-cultured with CS extract-exposed Tregs without or with bafilomycin. **(C)** Corrected total cell fluorescence (CTCF) was calculated using the FIJI Software, which calculates the integrated density of an image selected (in this case a MDM). The area of the MDM is multiplied by the mean fluorescence of the background image (an area of the image around the MDM). This number is then subtracted from the integrated density previously calculated. The equation was contributed by The QBI Advanced Microscopy facility, The University of Queensland, Australia.

**Supplementary Figure 3.**
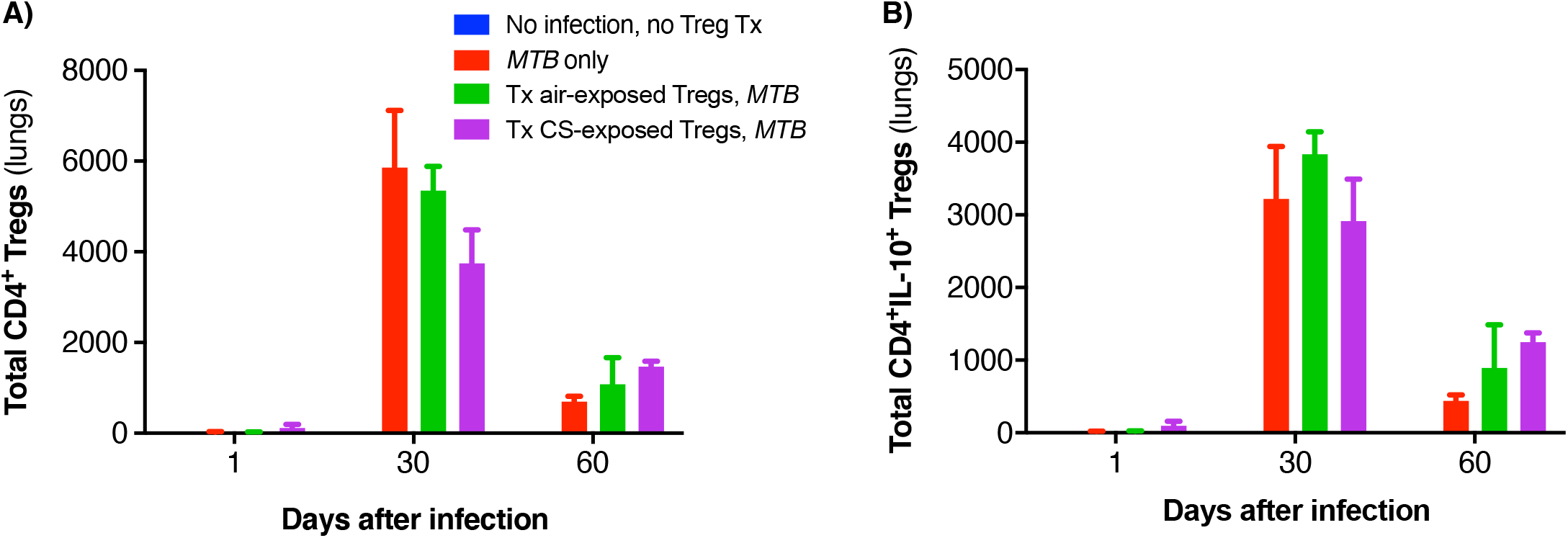
Quantitation of T regulatory cells from the lungs of mice that underwent adoptive transfer. **(A)** Total CD4^+^CD25^hi^CD127^-^ (T regulatory cells) were quantified in the lungs of uninfected mice, *Mycobacterium tuberculosis* (*MTB*)-infected mice, and Thy1.2 mice that received T regulatory cells (Tregs) from air- or cigarette smoke (CS)-exposed Thy1.1 mice and then infected with *MTB*. **(B)** Total CD4^+^CD25^hi^CD127^-^IL-10^+^ in the lungs of of the same mouse groups were quantified.

**Supplementary Figure 4.**
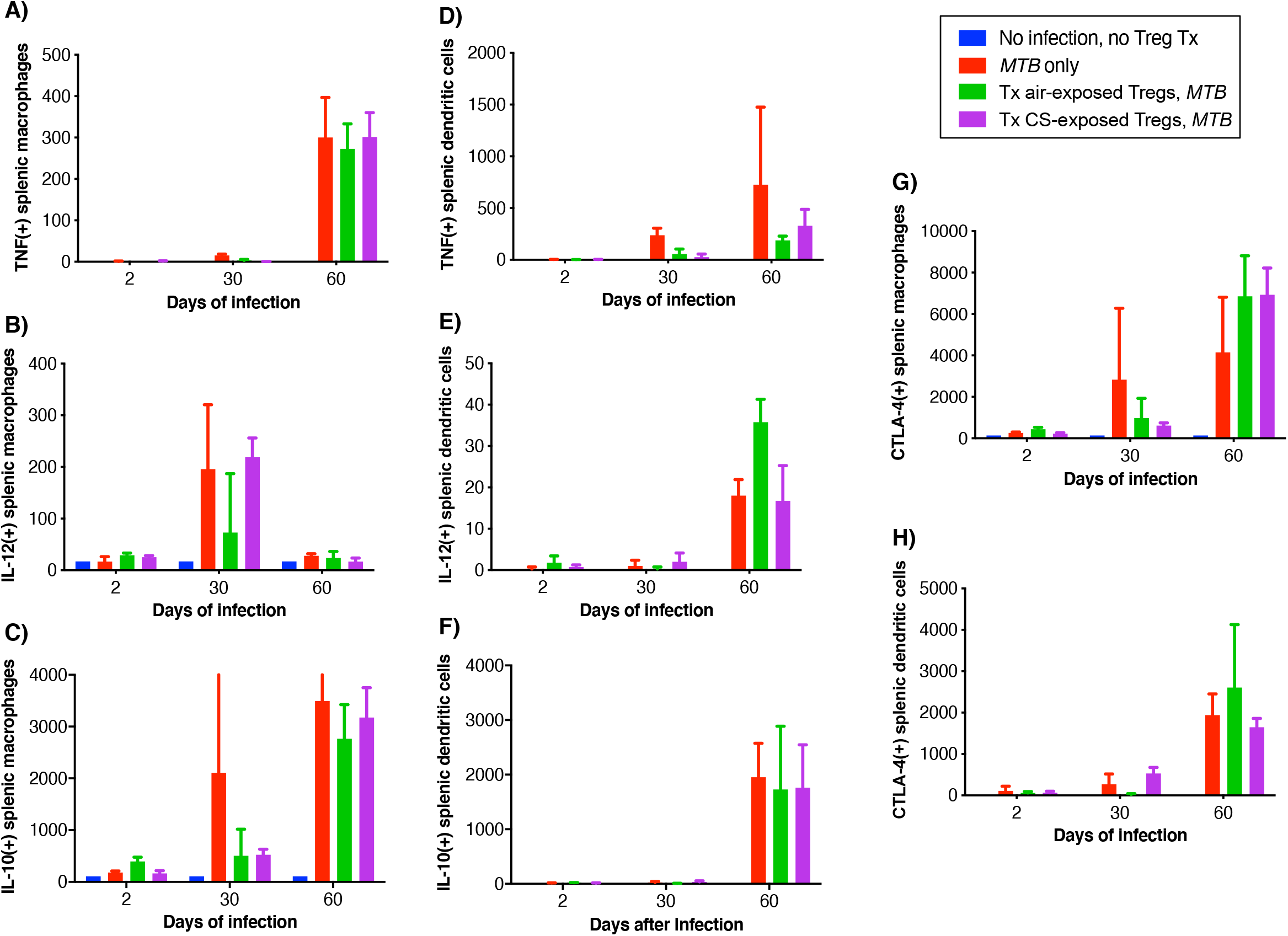
Intracellular cytokine and CTLA-4 analyses of murine spleen macrophages and dendritic cells. Splenic macrophages and dendritic cells (DC) from uninfected mice, *Mycobacterium tuberculosis* (*MTB*)-infected mice, and *MTB*-infected mice with adoptive transfer of Tregs from either air-exposed or CS-exposed mice were stained for **(A/D)** TNF, **(B/E)** IL-12, and **(C/F)** IL-10. From the same mouse groups, splenic macrophages and DC were stained for cell surface CTLA-4 **(G/H)**, respectively.

**Supplementary Figure 5.**
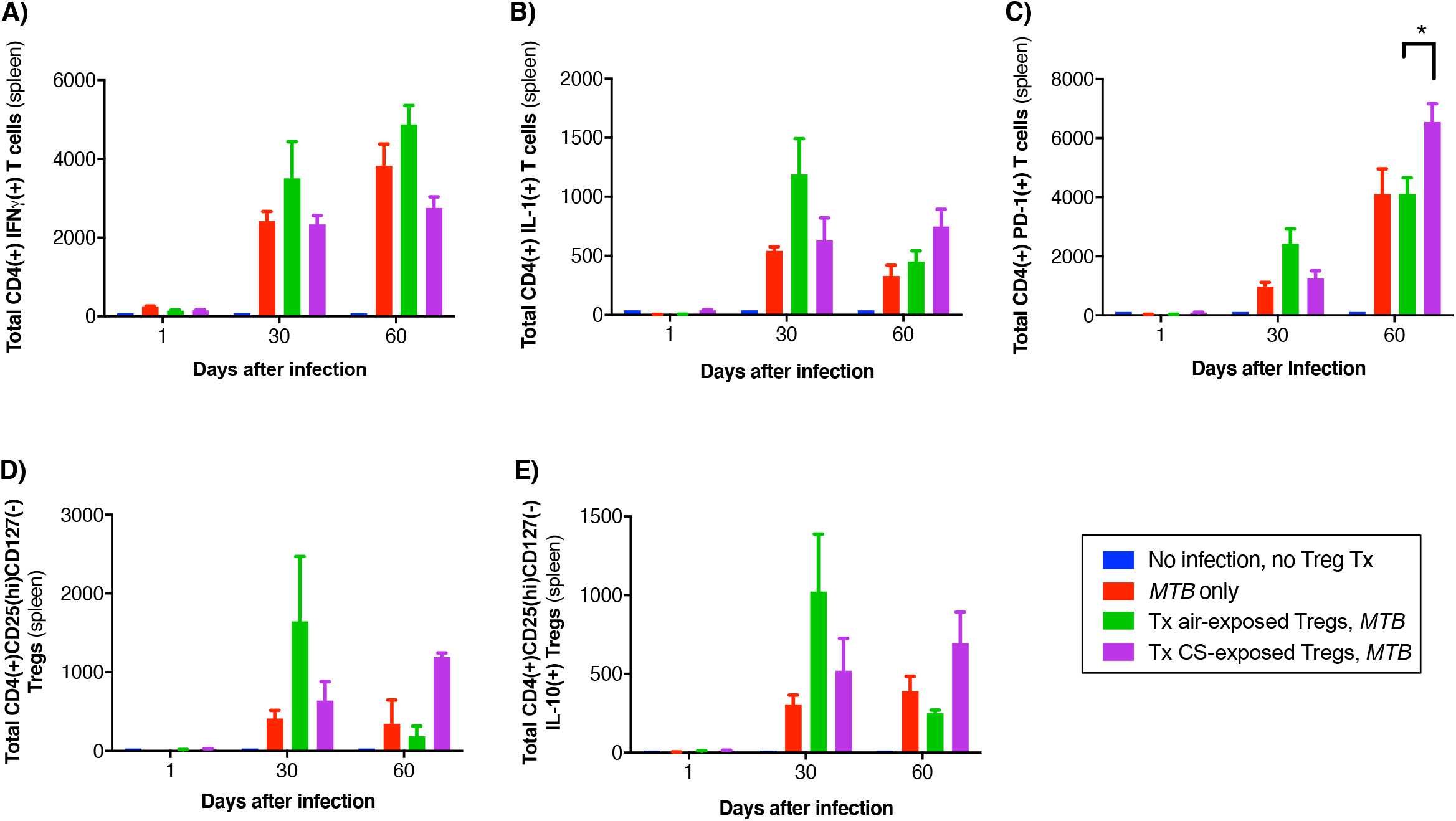
Intracellular cytokine and PD-1 analyses of murine splenic T cells. Splenic CD4^+^ T cells from uninfected mice, *Mycobacterium tuberculosis* (*MTB*)-infected mice, and *MTB*-infected mice with adoptive transfer of Tregulatory cells (Tregs) from either air-exposed or CS-exposed mice were stained for **(A)** IFN*γ*, **(B)** IL-1, and **(C)** PD-1 at the indicated time points after *MTB* infection. From the same mouse groups, **(D)** total splenic CD4^+^CD25^hi^CD127^-^ Tregs and **(E)** CD4^+^IL-10^+^ Tregs were quantified at the indicated times.

## References

1. Pai M, Mohan A, Dheda K, Leung CC, Yew WW, Christopher DJ, Sharma SK. 2007. Lethal interaction: the colliding epidemics of tobacco and tuberculosis. Expert Rev Anti Infect Ther 5:385–391.

2. Bishwakarma R, Kinney WH, Honda JR, Mya J, Strand MJ, Gangavelli A, Bai X, Ordway DJ, Iseman MD, Chan ED. 2015. Epidemiologic link between tuberculosis and cigarette/biomass smoke exposure: Limitations despite the vast literature. Respirology 20:556–568.

3. Chiang C-Y, Bam TS. 2020. The impact of smoking on TB treatment outcomes includes recurrent TB. Int J Tuberc Lung Dis 24:1224–1225.

4. Wang EY, Arrazola RA, Mathema B, Ahluwalia IB, Mase SR. 2020. The impact of smoking on tuberculosis treatment outcomes: a meta-analysis. Int J Tuberc Lung Dis 24:170–175.

5. Chan ED, Kinney WH, Honda JR, Bishwakarma R, Gangavelli A, Mya J, Bai X, Ordway DJ. 2014. Tobacco exposure and susceptibility to tuberculosis: is there a smoking gun? Tuberculosis (Edinb) 94:544–550.

6. Feng Y, Kong Y, Barnes PF, Huang FF, Klucar P, Wang X, Samten B, Sengupta M, Machona B, Donis R, Tvinnereim AR, Shams H. 2011. Exposure to cigarette smoke inhibits the pulmonary T-cell response to influenza virus and *Mycobacterium tuberculosis*. Infect Immun 79:229–237.

7. OLeary SM, Coleman MM, Chew WM, Morrow C, McLaughlin AM, Gleeson LE, OSullivan MP, Keane J. 2014. Cigarette smoking impairs human pulmonary immunity to *Mycobacterium tuberculosis*. Am J Respir Crit Care Med 190:1430–1436.

8. Shaler CR, Horvath CN, McCormick S, Jeyanathan M, Khera A, Zganiacz A, Kasinska J, Stampfli MR, Xing Z. 2013. Continuous and discontinuous cigarette smoke exposure differentially affects protective Th1 immunity against pulmonary tuberculosis. PLoS One 8:e59185.

9. Shang S, Ordway D, Henao-Tamayo M, Bai X, Oberley-Deegan R, Shanley C, Orme IM, Case S, Minor M, Ackart D, Hascall-Dove L, Ovrutsky AR, Kandasamy P, Voelker DR, Lambert C, Freed BM, Iseman MD, Basaraba RJ, Chan ED. 2011. Cigarette smoke increases susceptibility to tuberculosis – evidence from *in vivo* and *in vitro* models. J Infect Dis 203:1240–1248.

10. van Zyl-Smit RN, Binder A, Meldau R, Semple PL, Evans A, Smith P, Bateman ED, Dheda K. 2014. Cigarette smoke impairs cytokine responses and BCG containment in alveolar macrophages. Thorax 69:363–370.

11. Bai X, Stitzel JA, Bai A, Zambrano CA, Phillips M, Marrack P, Chan ED. 2017. Nicotine impairs macrophage control of *Mycobacterium tuberculosis*. Am J Respir Cell Mol Biol 57:324–333.

12. Ozeki Y, Sugawara I, Udagawa T, Aoki T, Osada-Oka M, Tateishi Y, Hisaeda H, Nishiuchi Y, Harada N, Kobayashi K, Matsumoto S. 2010. Transient role of CD4+CD25+regulatory T cells in mycobacterial infection in mice. Int Immunol 22:179–189.

13. Boer MC, Joosten SA, Ottenhoff TH. 2015. Regulatory T-Cells at the Interface between Human Host and Pathogens in Infectious Diseases and Vaccination. Front Immunol 6:217.

14. Larson RP, Shafiani S, Urdahl KB. 2013. Foxp3+regulatory T cells in tuberculosis, p 165–180. In Divangahi M (ed), The New Paradigm of Immunity to Tuberculosis. Springer, New York.

15. Ordway D, Henao-Tamayo M, Harton M, Palanisamy G, Troudt J, Shanley C, Basaraba RJ, Orme IM. 2007. The hypervirulent *Mycobacterium tuberculosis* strain HN878 induces a potent TH1 response followed by rapid down-regulation. J Immunol 179:522–531.

16. Rowe JH, Ertelt JM, Way SS. 2012. Foxp3(+) regulatory T cells, immune stimulation and host defence against infection. Immunology 136:1–10.

17. Semple PL, Binder AB, Davids M, Maredza A, van Zyl-Smit RN, Dheda K. 2013. Regulatory T-cells attenuate mycobacterial stasis in alveolar and blood-derived macrophages from patients with TB. Am J Respir Crit Care Med 187:1249–1258.

18. Shafiani S, Tucker-Heard G, Kariyone A, Takatsu K, Urdahl KB. 2010. Pathogen-specific regulatory T cells delay the arrival of effector T cells in the lung during early tuberculosis. J Exp Med 207:1409–1420.

19. Shang S, Harton M, Tamayo MH, Shanley C, Palansamy GS, Caraway M, Chan ED, Basaraba RJ, Orme IM, Ordway DJ. 2011. Increased Foxp3 expression in guinea pigs infected with W-Beijing strains of *M. tuberculosis*. Tuberculosis 91:378–385.

20. Barceló B, Pons J, Ferrer JM, Sauleda J, Fuster A, Agustí AG. 2008. Phenotypic characterisation of T-lymphocytes in COPD: abnormal CD4+CD25+regulatory T-lymphocyte response to tobacco smoking. Eur Respir J 31:555–562.

21. Botelho FM, Gaschler GJ, Kianpout S, Zavitz CCJ, Trimble NJ, Nikota JK, Bauer AMT, Stampfli MR. 2010. Innate immune processes are sufficient for driving cigarette smoke-induced inflammation in mice. Am J Respir Cell Mol Biol 42:394–403.

22. Ménard L, Rola-Pleszczynski M. 1987. Nicotine induces T-suppressor cells: modulation by the nicotinic antagonist D-tubocurarine and myasthenic serum. Clin Immunol Immunopathol 44:107–113.

23. Wang DW, Zhou RB, Yao YM, Zhu XM, Yin YM, Zhao GJ, Dong N, Sheng ZY. 2010. Stimulation of α7 nicotinic acetylcholine receptor by nicotine increases suppressive capacity of naturally occurring CD4+CD25+regulatory T cells in mice *in vitro*. J Pharmacol Exp Ther 335:553–561.

24. Wallace WA, Gillooly M, Lamb D. 1992. Intra-alveolar macrophage numbers in current smokers and non-smokers: a morphometric study of tissue sections. Thorax 47:437–440.

25. Ouyang Y, Virasch N, Hao P, Aubrey MT, Mukerjee N, Bierer BE, Freed BM. 2000. Suppression of human IL-1beta, IL-2, IFN-gamma, and TNF-alpha production by cigarette smoke extracts. J Allergy Clin Immunol 106:280–287.

26. Bai X, Shang S, Henao-Tamayo M, Basaraba RJ, Ovrutsky AR, Matsuda JL, Takeda K, Chan MM, Dakhama A, Kinney WH, Trostel J, Bai A, Honda JR, Achcar R, Hartney J, Joosten LAB, Kim S-H, Orme IM, Dinarello CA, Ordway DJ, Chan ED. 2015. Human IL-32 expression protects mice against a hypervirulent strain of *Mycobacterium tuberculosis*. Proc Natl Acad Sci USA 112:5111–5116.

27. Ko JH, Yoon SO, Lee HJ, Oh JY. 2017. Rapamycin regulates macrophage activation by inhibiting NLRP3 inflammasome-p38 MAPK-NFκB pathways in autophagy- and p62-dependent manners. Oncotarget 8:40817–40831.

28. Mauvezin C, Neufeld TP. 2015. Bafilomycin A1 Disrupts Autophagic Flux by Inhibiting Both V–ATPase–dependent Acidification and Ca–P60A/SERCA–dependent Autophagosome–Lysosome Fusion. Autophagy 11:1437–1438.

29. Hammond L. 2014. Measuring cell fluorescence using ImageJ. https://theolb.readthedocs.io/en/latest/imaging/measuring-cell-fluorescence-using-imagej.html. Accessed

30. Francisco LM, Sage PT, Sharpe AH. 2010. The PD-1 pathway in tolerance and autoimmunity. Immunol Rev 236:219–242.

31. Yanagita M, Kobayashi R, Kojima Y, Mori K, Murakami S. 2012. Nicotine modulates the immunological function of dendritic cells through peroxisome proliferator-activated receptor-*γ* upregulation. Cell Immunol 274:26–33.

32. Saresella M, Marventano I, Longhi R, Lissoni F, Trabattoni D, Mendozzi L, Caputo D, Clerici M. 2008. CD4+CD25+FoxP3+PD1-regulatory T cells in acute and stable relapsing-remitting multiple sclerosis and their modulation by therapy. FASEB J 22:3500–3508.

33. Machicote A, Belén S, Baz P, Billordo LA, Fainboim L. 2018. Human CD8+HLA-DR+Regulatory T Cells, Similarly to Classical CD4+Foxp3+Cells, Suppress Immune Responses via PD-1 / PD-L1 Axis. Front Immunol 9:2788.

34. Laurent S, Carrega P, Saverino D, Piccioli P, Camoriano M, Morabito A, Dozin B, Fontana V, Simone R, Mortara L, Mingari MC, Ferlazzo G, Pistillo MP. 2010. CTLA-4 is expressed by human monocyte-derived dendritic cells and regulates their functions. Hum Immunol 71:934–941.

35. Koenecke C, Lee CW, Thamm K, Föhse L, Schafferus M, Mittrücker HW, Floess S, Huehn J, Ganser A, Förster R, Prinz I. 2012. IFN-*γ* production by allogeneic Foxp3+regulatory T cells is essential for preventing experimental graft-versus-host disease. J Immunol 189:2890–2896.

36. Zhao J, Zhao J, Fett C, Trandem K, Fleming E, Perlman S. 2011. IFN-*γ*- and IL-10-expressing virus epitope-specific Foxp3(+) T reg cells in the central nervous system during encephalomyelitis. J Exp Med 208:1571–1577.

37. Sumida T, Lincoln MR, Ukeje CM, Rodriguez DM, Akazawa H, Noda T, Naito AT, Komuro I, Dominguez-Villar M, Hafler DA. 2018. Activated beta-catenin in Foxp3(+) regulatory T cells links inflammatory environments to autoimmunity. Nat Immunol 19:1391–1402.

38. Henao-Tamayo MI, Obregón-Henao A, Arnett K, Shanley CA, Podell B, Orme IM, Ordway DJ. 2016. Effect of Bacillus Calmette-Guérin Vaccination on CD4+Foxp3+T Cells During Acquired Immune Response to *Mycobacterium tuberculosis* Infection. J Leukoc Biol 99:605–617.

39. Shaykhiev R, Krause A, Salit J, Strulovici-Barel Y, Harvey BG, OConnor TP, Crystal RG. 2009. Smoking-dependent reprogramming of alveolar macrophage polarization: implication for pathogenesis of chronic obstructive pulmonary disease. J Immunol 183:2867–2883.

40. Cavassani KA, Carson WF, Moreira AP, Wen H, Schaller MA, Ishii M, Lindell DM, Dou Y, Lukacs NW, Keshamouni VG, Hogaboam CM, Kunkel SL. 2010. The post sepsis-induced expansion and enhanced function of regulatory T cells create an environment to potentiate tumor growth. Blood 115:4403–4411.

41. Shevach EM. 2009. Mechanisms of Foxp3+T regulatory cell-mediated suppression. Immunity 30:636–645.

42. Vignali DA, Collison LW, Workman CJ. 2008. How regulatory T cells work. Nat Rev Immunol 8:523–532.

43. Grossman WJ, Verbsky JW, Barchet W, Colonna M, Atkinson JP, Ley TJ. 2004. Human T regulatory cells can use the perforin pathway to cause autologous target cell death. Immunity 21:589–601.

44. Martinez FO, Helming L, Gordon S. 2009. Alternative activation of macrophages: an immunologic functional perspective. Annu Rev Immunol 27:451–483.

45. Tiemessen MM, Jagger AL, Evans HG, van Herwijnen MJ, John S, Taams LS. 2007. CD4+CD25+Foxp3+regulatory T cells induce alternative activation of human monocytes/macrophages. Proc Natl Acad Sci U S A 104:19446–19451.

46. Mills CD. 2012. M1 and M2 Macrophages: Oracles of Health and Disease. Crit Rev Immunol 32:463–488.

47. Romano M, Fanelli G, Tan N, Nova-Lamperti E, McGregor R, Lechler RI, Lombardi G, Scottà C. 2018. Expanded Regulatory T Cells Induce Alternatively Activated Monocytes With a Reduced Capacity to Expand T Helper-17 Cells. Front Immunol 9:1625.

48. Liang B, Workman C, Lee J, Chew C, Dale BM, Colonna L, Flores M, Li N, Schweighoffer E, Greenberg S, Tybulewicz V, Vignali D, Clynes R. 2008. Regulatory T cells inhibit dendritic cells by lymphocyte activation gene-3 engagement of MHC class II. J Immunol 180:5916–5926.

49. Yu X, Harden K, Gonzalez LC, Francesco M, Chiang E, Irving B, Tom I, Ivelja S, Refino CJ, Clark H, Eaton D, Grogan JL. 2009. The surface protein TIGIT suppresses T cell activation by promoting the generation of mature immunoregulatory dendritic cells. Nat Immunol 10:48–57.

50. Cardona P, Cardona P-J. 2019. Regulatory T Cells in *Mycobacterium tuberculosis* Infection. Front Immunol 10:2139.

51. Kumar P. 2017. IFN*γ*-producing CD4 +T lymphocytes: the double-edged swords in tuberculosis. Clin Transl Med 6:21.

52. Zhang S, Petro TM. 1996. The effect of nicotine on murine CD4 T cell responses. Int J Immunopharmacol 18:467–478.

53. Wing K, Onishi Y, Prieto-Martin P, Yamaguchi T, Miyara M, Fehervari Z, Nomura T, Sakaguchi S. 2008. CTLA-4 control over Foxp3+regulatory T cell function. Science 322:271–275.

54. Walker LS. 2013. Treg and CTLA-4: two intertwining pathways to immune tolerance. J Autoimmun 45:49–57.

55. Kong YM, Flynn JC. 2014. Opportunistic autoimmune disorders potentiated by immune-checkpoint inhibitors anti-CTLA-4 and anti-PD-1. Front Immunol 5:206.

56. Hua GY, Wang P, Takagi K, Shimozato O, Yagita H, Okigaki T, Matasumura M. 1999. Expression of a soluble form of CTLA4 on macrophage and its biological activity. Cell Res 9:189–199.

57. Wang XB, Giscombe R, Yan Z, Heiden T, Xu D, Lefvert AK. 2002. Expression of CTLA-4 by human monocytes. Scand J Immunol 55:53–60.

58. Wang XB, Fan ZZ, Anton D, Vollenhoven AV, Ni ZH, Chen XF, Lefvert AK. 2011. CTLA4 is expressed on mature dendritic cells derived from human monocytes and influences their maturation and antigen presentation. BMC Immunol 12:21.

